# The ESCRT protein CHMP5 restricts bone formation by controlling endolysosome-mitochondrion-mediated cell senescence

**DOI:** 10.1101/2020.08.03.233874

**Authors:** Fan Zhang, Yuan Wang, Luyang Zhang, Chunjie Wang, Deping Chen, Haibo Liu, Ren Xu, Cole M. Haynes, Jae-Hyuck Shim, Xianpeng Ge

## Abstract

The dysfunction of the cellular endolysosomal pathway, such as in lysosomal storage diseases, can cause severe musculoskeletal disorders. However, how endolysosomal dysfunction causes musculoskeletal abnormalities remains poorly understood, limiting therapeutic options. Here, we report that CHMP5, a member of the endosomal sorting complex required for transport (ESCRT)-III protein family, is essential to maintain the endolysosomal pathway and regulate bone formation in osteogenic lineage cells. Genetic ablation of *Chmp5* in mouse osteogenic cells increases bone formation in vivo and in vitro. Mechanistically, *Chmp5* deletion causes endolysosomal dysfunction by decreasing the VPS4A protein, and CHMP5 overexpression is sufficient to increase the VPS4A protein. Subsequently, endolysosomal dysfunction disturbs mitochondrial functions and increases mitochondrial ROS, ultimately resulting in skeletal cell senescence. Senescent skeletal cells cause abnormal bone formation by combining cell-autonomous and paracrine actions. Importantly, the elimination of senescent cells using senolytic drugs can alleviate musculoskeletal abnormalities in *Chmp5* conditional knockout mice. Therefore, our results show that cell senescence represents an underpinning mechanism and a therapeutic target for musculoskeletal disorders caused by the aberrant endolysosomal pathway, such as in lysosomal storage diseases. These results also uncover the function and mechanism of CHMP5 in the regulation of cell senescence by affecting the endolysosomal-mitochondrial pathway.

## Introduction

The endosomal sorting complex required for transport (ESCRT) machinery is an evolutionarily conserved molecular mechanism essential for diverse cell biological processes. Since it was initially discovered in the early 21st century in the budding yeast *Saccharomyces cerevisiae*, the molecular gear has been extensively dissected in the endocytic pathway, which sorts ubiquitinated membrane proteins and bound cargoes that commit to lysosomal degradation [1–4]. In addition to the endocytic process, ESCRT proteins also play critical roles in other cell activities, such as exocytosis, viral budding, cytokinesis, pruning of neurons, repair of the plasma membrane, and assembly of the nuclear envelope [2, 4]. In the endocytic pathway, the ESCRT machinery consists of a series of functionally overlapping protein complexes, including ESCRT-0, -I, -II and -III. While the ESCRT-0, -I, and -II complexes are mainly responsible for protein sorting and recruiting ESCRT-III components, the ESCRT-III complex, cooperating with the ATPase VPS4, is the primary executor of membrane severing during the formation of multivesicular bodies (MVB) [3, 5], which ultimately fuse with the lysosomes for cargo degradation or with the plasma membrane to release cargos into the extracellular space.

Charged multivesicular body protein 5 (CHMP5), the mammalian ortholog of yeast VPS60/MOS10, is a component of the ESCRT-III protein complex and plays an essential role in the late stage of MVB formation [6]. As the yeast *Vps60/Mos10*-null mutation disturbs the endosome-to-vacuole (the lysosome-like structure in yeast) trafficking and affects the sorting of internalized endocytic markers [7], ablation of *Chmp5* in mouse embryonic fibroblasts (MEFs) also affects the sorting from late endosomes/MVBs to lysosomes and reduces lysosomal degradation of multiple activated receptors [6]. Notably, in mouse osteoclasts, CHMP5 suppresses ubiquitylation of the IκBα protein and restricts the activity of NFκB pathway, without any effect on endolysosomal functions [8]. During αβ T lymphocyte development, CHMP5 promotes thymocyte survival by stabilizing the pro-survival protein of BCL2 and does not have an influence on the endolysosomal system [9]. These studies suggest that the regulation of the endolysosomal pathway by CHMP5 depends on the cell and tissue context.

The dysfunction of the endolysosomal pathway can cause severe musculoskeletal disorders. A such example is lysosomal storge diseases, which include more than 50 different subgroups of diseases due to mutations in genes that encode lysosomal hydrolases or genes that are responsible for the transport, maturation, and functions of lysosomal hydrolases [10]. Musculoskeletal pathologies, such as joint stiffness/contracture, bone deformation, muscular atrophy, short stature, and decreased mobility, are among the most common, in some cases the earliest, clinical presentations of lysosomal storage diseases [11, 12]. These musculoskeletal lesions are particularly refractory to current treatments for lysosomal storage diseases, including enzyme replacement treatment and hematopoietic stem cell transplantation [13]. Therefore, studies on the mechanisms of musculoskeletal pathologies due to endolysosomal dysfunction are paramount to design and develop novel treatments for these disorders.

In this study, our results uncover the function and mechanism of CHMP5 in the regulation of cell senescence and bone formation in osteogenic cells. Deletion of *Chmp5* causes endolysosomal dysfunction involving decreased VPS4A protein and subsequently activates cell senescence by increasing intracellular mitochondrial ROS. Senescent cells induce bone formation in both autonomous and paracrine ways. Importantly, senolytic treatment is effective in mitigating musculoskeletal pathologies caused by *Chmp5* deletion.

## Results

### Ablation of *Chmp5* in mouse osteogenic lineage cells causes aberrant bone formation

In addition to the well-characterized role of the CTSK gene in osteoclasts, this gene has recently been reported to identify periskeletal progenitors [14–16]. In particular, the ablation of *Chmp5* in mouse *Ctsk*-expressing cells (hereafter referred to as *Chmp5^Ctsk^* mice, homozygous) caused dramatic periskeletal bone overgrowth near the joint compared to *Chmp5^Ctsk/+^* (heterozygous) and *Chmp5^fl/fl^* littermate controls (**Fig. 1A and B**). Aberrant periskeletal bone growth in *Chmp5^Ctsk^* mice became distinct around postnatal day 10 (d10), progressed with age, and involved multiple bones and joints throughout the body, including knee, ankle, and foot joints (**Fig. 1A to C and Fig. S1A**). With age, these animals developed severe skeletal deformities, progressive joint stiffness, reduced motility, and short stature. Despite bone overgrowth in *Chmp5^Ctsk^* animals, mechanical tests showed that bone stiffness and fracture stress decreased significantly (**Fig. 1D**), suggesting that bone mechanical properties were impaired in *Chmp5^Ctsk^* mice.

**Fig. 1.**
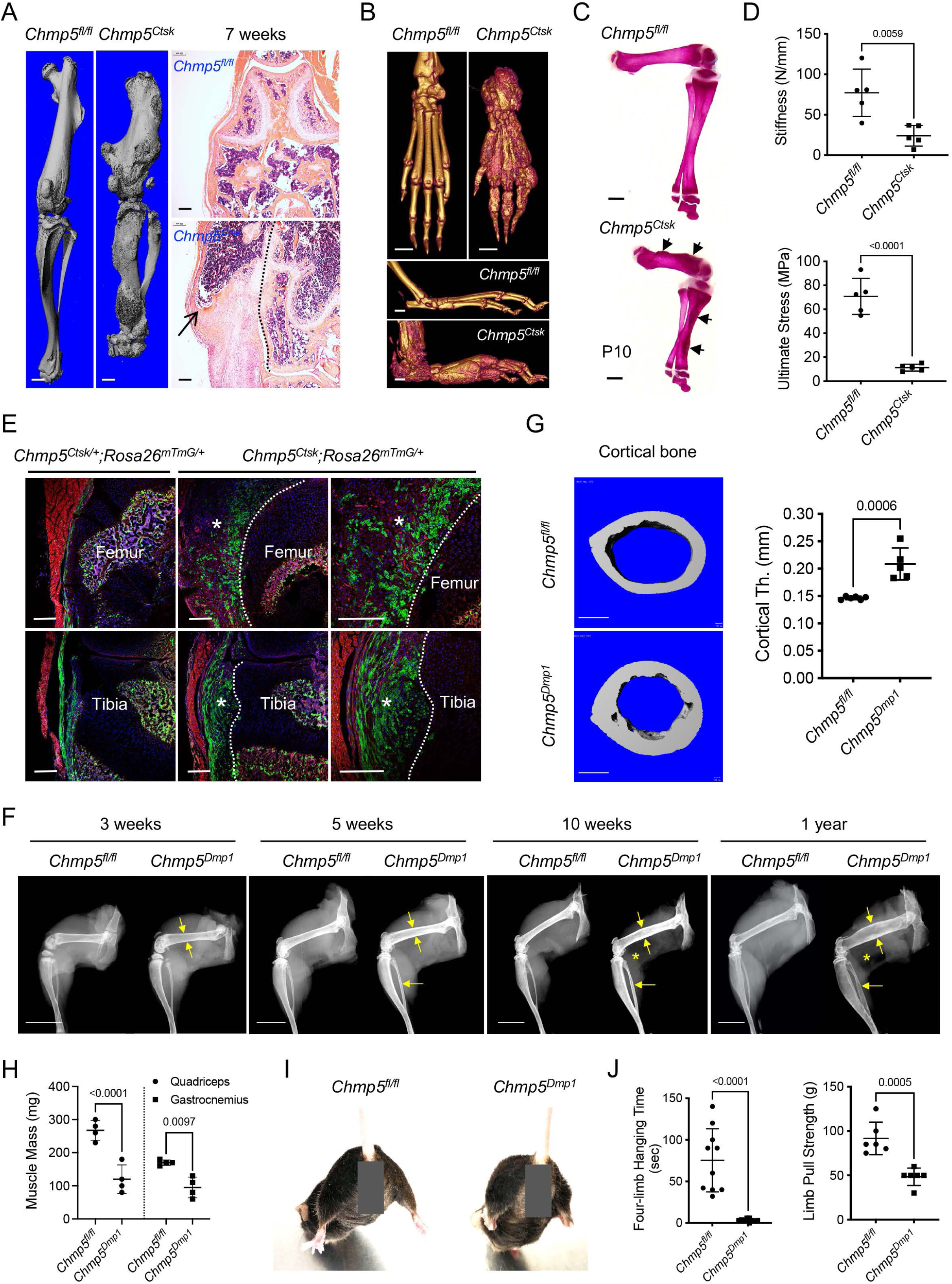
Ablation of *Chmp5* in mouse osteogenic lineage cells causes aberrant bone formation. (**A**) Micro-CT and H&E staining showing periskeletal overgrowth in *Chmp5^Ctsk^* mice in comparison with *Chmp5^fl/fl^* mice at 7 weeks of age. Dot line representing the approximate bone border. *n* = 10 animals per group. (**B**) Micro-CT images displaying periskeletal overgrowth in the ankle and foot in *Chmp5^Ctsk^* versus *Chmp5^fl/fl^* mice at one year of age. *n* = 4 animals per group. (**C**) Alizarin red staining of skeletal preparations on postnatal day 10. *n* = 4 mice per group, arrows indicating periskeletal bone overgrowth. (**D**) Three-point test showing femur bone stiffness and ultimate stress (fracture stress) in *Chmp5^Ctsk^* and *Chmp5^fl/fl^* mice. *n* = 5 male animals per group. Similar changes observed in both sexes. (**E**) Confocal images mapping *Ctsk*^+^ (GFP^+^) progenitors in periskeletal tissues in *Chmp5^Ctsk/+^;Rosa26^mTmG/+^* and *Chmp5^Ctsk^;Rosa26^mTmG/+^* mice at 2 weeks of age. Asterisks indicating periskeletal overgrowths; dot-line representing the approximate bone border. *n* = 4 animals each group. (**F**) X-ray images demonstrating progressive cortical bone expansion (arrows) and decreased skeletal muscle mass (asterisks) in *Chmp5^Dmp1^* in comparison with *Chmp5^fl/fl^* mice. *n* = 3, 3, 10, 8 animals per group for 3 weeks, 5 weeks, 10 weeks, and 1 year of age, respectively. (**G**) Micro-CT analyses displaying cortical bone expansion in *Chmp5^Dmp1^* relative to *Chmp5^fl/fl^* mice at 10 weeks of age. *n* = 6 *Chmp5^fl/fl^* and 5 *Chmp5^Dmp1^* male mice, similar changes found in both genders. (**H**) Skeletal muscle mass in *Chmp5^Dmp1^* compared to *Chmp5^fl/fl^* mice at 10-12 weeks of age. *n* = 4 animals (male) per group, similar changes found in both genders. (**I**) Hindlimb abduction of *Chmp5^Dmp1^* mice in comparison with *Chmp5^fl/fl^* littermate controls at 10 weeks of age. More than 20 animals per group observed. (**J**) Four-limb handing time (*n* = 10 per group, pooled data from both sexes), and forelimb pull strength (*n* = 6 per group, male mice) in *Chmp5^Dmp1^* compared to *Chmp5^fl/fl^* mice at 10-12 weeks of age. Similar changes found in both genders. All data are mean ± s.d.; two-tailed unpaired Student’s *t*-test. Scale bars, 1 mm for micro-CT and skeletal preparation images, 200 μm for histological images, and 10 mm in panel (**F**).

To track CTSK^+^ periskeletal progenitors during the bone overgrowth in *Chmp5^Ctsk^* mice, we crossed *Ctsk-Cre*, *Chmp5^fl/fl^*, and *Rosa26^mTmG^* mice to generate *Chmp5^Ctsk/+^;Rosa26^mTmG/+^* and *Chmp5^Ctsk^;Rosa26^mTmG/+^* reporter animals, in which CTSK^+^ periskeletal progenitors and their descendants express membrane-localized green fluorescence protein (GFP). In *Chmp5^Ctsk/+^;Rosa26^mTmG/+^* control mice, GFP^+^ cells could be detected in the perichondrium, groove of Ranvier, periosteum, ligament, and enthesis in addition to osteoclasts in the bone marrow, but not in growth plate chondrocytes (**Fig. 1E**). With *Chmp5* deletion, GFP^+^ cells consisted of a major part of cells in periskeletal overgrowth in *Chmp5^Ctsk^;Rosa26^mTmG/+^* mice (**Fig. 1E** and **Fig. S1B**). Tartarate resistant acid phosphatase (TRAP) staining did not show TRAP^+^ osteoclasts in the periskeletal lesion, while obvious TRAP^+^ osteoclasts could be detected in the bone marrow (**Fig. S1C**). These results indicate that *Chmp5*-deficient periskeletal progenitors directly contribute to the bone overgrowth in *Chmp5^Ctsk^* mice.

Next, we sought to confirm the role of CHMP5 in the regulation of bone formation by deleting this gene using another mouse line of osteogenic lineage Cre. We crossed *Chmp5^fl/fl^* mice with the line of *Prrx1*-Cre [17], *Col2*-Cre [18] or *Osx*-Cre [19] mice to delete the *Chmp5* gene in *Prrx1*-expressing limb mesenchymal progenitors (*Chmp5^Prrx1^* mice), *Col2*-expressing perichondral progenitors (*Chmp5^Col2^* mice) or *Osx*-expressing osteoprogenitors (*Chmp5^Osx^* mice) and were unable to obtain postnatal homozygous gene knockout mice, probably due to embryonic lethality after ablating C*hmp5* expression in these osteogenic lineages. Subsequently, we used the *Dmp1*-Cre mouse line that could target postnatal endosteal osteoprogenitors, preosteoblasts, mature osteoblasts, and osteocytes [20–22]. *Chmp5^Dmp1^* mice were born in normal Mendelian ratios and were viable. Within three weeks of age, the skeletons of *Chmp5^Dmp1^* mice began to show a distinct expansion in comparison with *Chmp5^Dmp1/+^* and *Chmp5^fl/fl^* littermate controls, which progressed with age and became much more severe at 1 year of age (**Fig. 1F**). Bone expansion in *Chmp5^Dmp1^* mice that occurred mainly in the endosteum was further shown in detail around the age of 10 weeks by micro-CT and histological analyses (**Fig. 1G** and **Fig. S1D, E**).

Meanwhile, since *Dmp1*-expressing lineage cells also generate skeletal muscle cells [21], *Chmp5^Dmp1^* mice showed a profound decrease in skeletal muscle mass with age (**Fig. 1F and H**). Hindlimb abduction, full-limb hanging time, and front-limb pull strength tests showed that muscular functions also decreased profoundly in *Chmp5^Dmp1^* compared to *Chmp5^fl/fl^* mice (**Fig. 1I and J**).

### CHMP5 restricts osteogenesis in skeletal progenitor cells

To further characterize the function of CHMP5 in the regulation of bone formation, we cultured and sorted *Ctsk*-expressing periskeletal progenitor cells (CD45^−^;CD31^−^;GFP^+^) from periskeletal tissues of *Chmp5^Ctsk/+^;Rosa26^mTmG/+^* and *Chmp5^Ctsk^;Rosa26^mTmG/+^* mice in the early stage of skeletal lesions (P10-P14) (**Fig. 2A and Fig. S2A, B**, hereafter referred to as *Chmp5^Ctsk/+^* and *Chmp5^Ctsk^* periskeletal progenitors, respectively). *Chmp5* deletion in *Chmp5^Ctsk^* periskeletal progenitors was confirmed by gene expression analysis (**Fig. 2B**). Strikingly, when cultured in the osteogenic differentiation medium, *Chmp5^Ctsk^* periskeletal progenitors showed considerably enhanced osteogenesis in comparison with *Chmp5^Ctsk/+^* control cells, as shown by alizarin red staining, von Kosaa staining, and alkaline phosphatase activity assay (**Fig. 2C**).

**Fig. 2.**
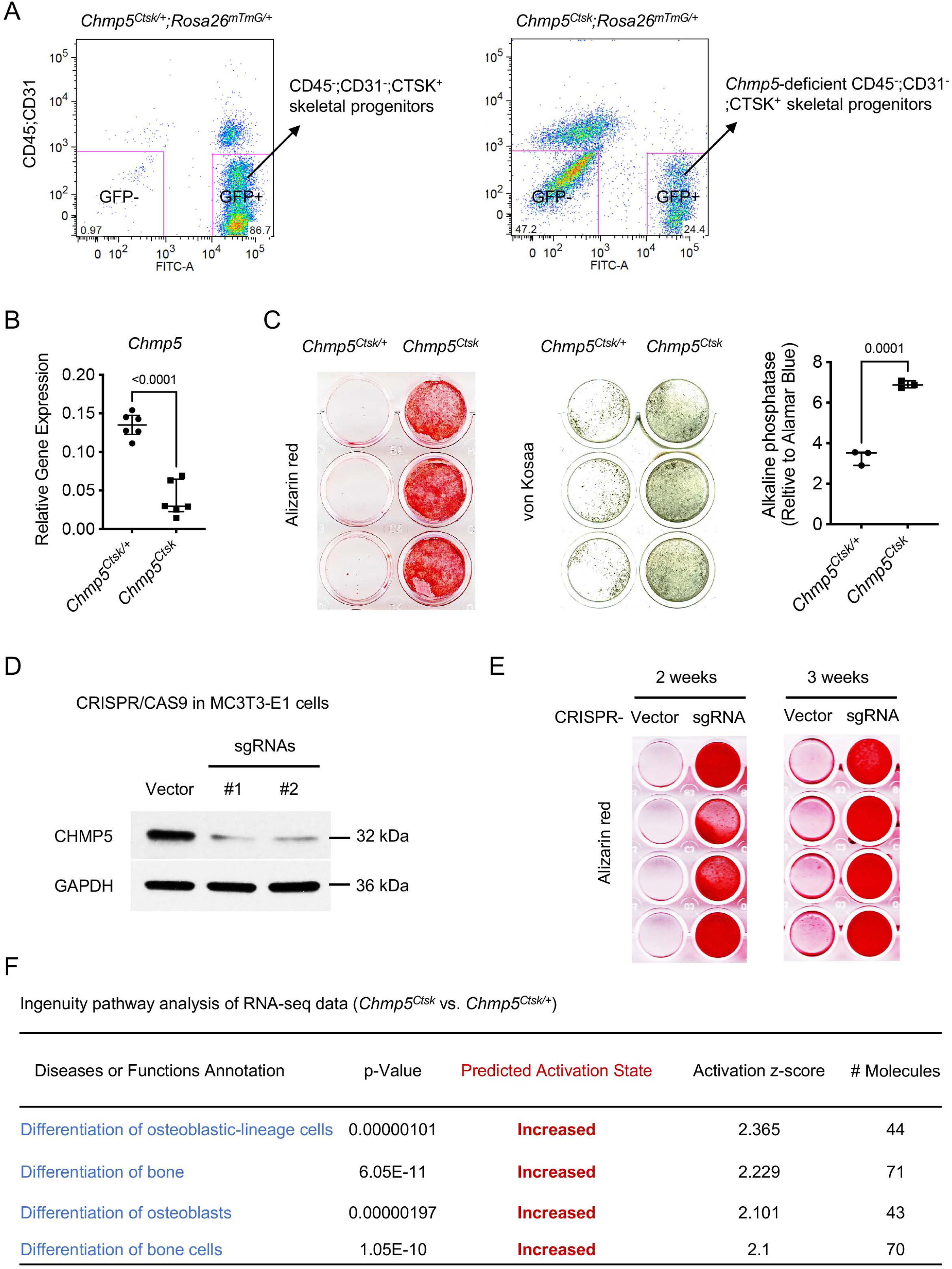
CHMP5 restricts osteogenesis in skeletal progenitor cells. (**A**) Flow cytometry showing the strategy for sorting CD45^−^;CD31^−^;GFP^+^ and CD45^−^;CD31^−^;GFP-periskeletal progenitors from periskeletal tissues of *Chmp5^Ctsk/+^;Rosa26^mTmG/+^* and *Chmp5^Ctsk^;Rosa26^mTmG/+^* mice at 2 weeks of age. *n* = 10 mice per group. (**B**) Quantitative PCR determining the expression of *Chmp5* in *Chmp5^Ctsk^* relative to *Chmp5^Ctsk/+^* periskeletal progenitors. *n* = 6 mice per group. (**C**) Alizarin red staining, von Kossa staining and alkaline phosphatase activity assay determining osteogenesis in *Chmp5^Ctsk^* compared to *Chmp5^Ctsk/+^* periskeletal progenitors. *n* = 3 replicates each group, representative results of cells from 3 different mice per group. (**D**) Western blot confirming *Chmp5* deletion in mouse MC3T3-E1 cells by lentiviral CRISPR/CAS9. (**E**) Alizarin red staining demonstrating osteogenesis in MC3T3-E1 cells with or without *Chmp5* deletion by lentiviral CRISPR/CAS9. *n* = 4 replicates per group, experiments repeated twice for each time point. **(F)** Ingenuity pathway analysis of RNA-seq data showing increased activity of osteoblast differentiation in *Chmp5^Ctsk^* vs. *Chmp5^Ctsk/+^* periskeletal progenitors. All data are mean ± s.d.; two-tailed unpaired Student’s *t*-test.

The regulation of osteogenesis by CHMP5 in skeletal progenitors was further determined in the osteogenic cell line MC3T3-E1 after removing *Chmp5* using CRISPR/CAS9 technology (**Fig. 2D**). The deletion of *Chmp5* in MC3T3-E1 cells also markedly increased osteogenesis, as shown by alizarin red staining at 2 weeks and 3 weeks of osteogenic induction (**Fig. 2E**). Consistently, the ingenuity pathway analysis of RNA-seq data of *Chmp5^Ctsk^* vs. *Chmp5^Ctsk/+^* periskeletal progenitors showed that multiple molecular pathways related to osteoblast differentiation were activated with the *Chmp5* deletion (**Fig. 2F**). Taken together, these results reveal a function of CHMP5 in regulating osteogenesis in skeletal progenitors.

### CHMP5 controls skeletal progenitor cell senescence

To gain more insight into the mechanism of regulation of bone formation by CHMP5, we assumed that depletion of *Chmp5* in osteogenic lineage cells also increases cell proliferation, which contributes to the bone overgrowth in *Chmp5* conditional knockout mice. Unexpectedly, *Chmp5^Ctsk^* periskeletal progenitors showed a significant decrease in the cell proliferation rate compared to *Chmp5^Ctsk/+^* controls (approximately 58% decrease in the cell number on day 6 of P2 culture, **Fig. 3A**). Because CHMP5 has been reported to regulate thymocyte apoptosis [9], we performed cell apoptosis analyses for *Chmp5^Ctsk^* and *Chmp5^Ctsk/+^* cells. Annexin V staining showed that approximately 4.96 ± 0.81% of *Chmp5^Ctsk^* periskeletal progenitors vs. 1.69 ± 0.68% of *Chmp5^Ctsk/+^* control cells were positively stained (**Fig. S3A**). Similarly, the in-situ terminal deoxynucleotidyl transferase dUTP nick end labelling assay (TUNEL) demonstrated that around 3.70 ± 0.72% cells in the periskeletal lesion of *Chmp5^Ctsk^* mice were positively labelled (**Fig. S3B**). These results show that cell apoptosis is activated in *Chmp5^Ctsk^* periskeletal progenitors in a rather low ratio.

**Fig. 3.**
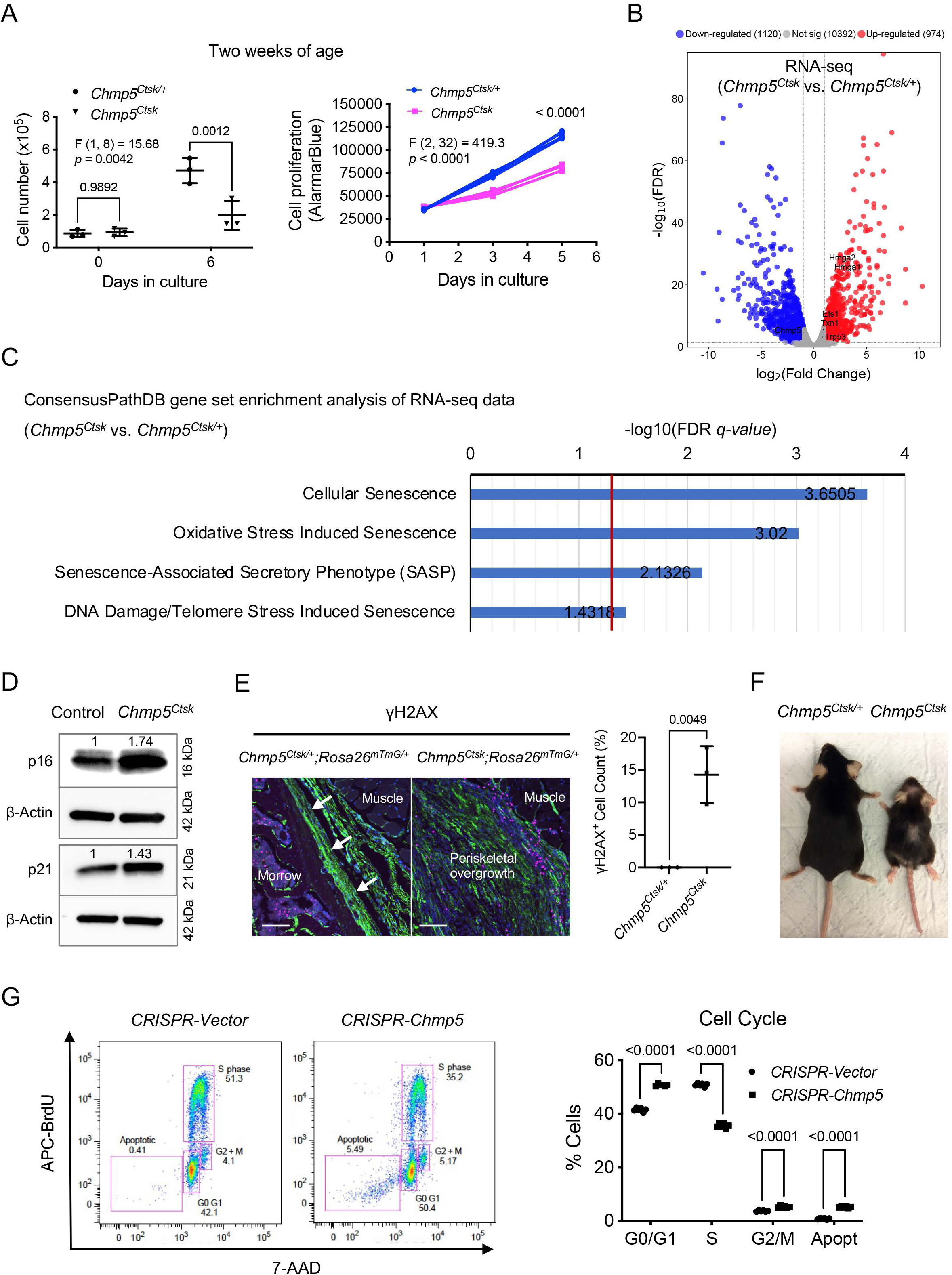
CHMP5 controls skeletal progenitor cell senescence. (**A**) Cell number counting and AlamarBlue assay determining cell proliferation in *Chmp5^Ctsk^* and *Chmp5^Ctsk/+^* periskeletal progenitors from 2-week-old animals. *n* = 3 replicates per group per time point; experiments repeated 3 times using cells from 3 different mice. (**B**) Volcano plot of RNA-seq data showing differentially expressed genes in *Chmp5^Ctsk^* vs. *Chmp5^Ctsk/+^* periskeletal progenitors. *n* = 3 per group with cells from 3 different animals. (**C**) ConsensusPathDB gene set enrichment analysis of RNA-seq data showing positive enrichment of genes in multiple molecular pathways related to cell senescence (Reactome) in *Chmp5^Ctsk^* relative to *Chmp5^Ctsk/+^* periskeletal progenitors. (**D**) Western blot determining the expression of p16 and p21 proteins in *Chmp5^Ctsk^* compared to *wild-type* (control) periskeletal progenitors. The numbers above lanes indicating the intensity of the p16 or p21 band relative to that of *Control* cells after normalization by β-Actin. Results repeated 3 times with cells from 3 different animals. (**E**) Immunostaining of γH2AX and quantification of γH2AX^+^;GFP^+^ cells in the periosteum and periskeletal overgrowth in *Chmp5^Ctsk/+^;Rosa26^mTmG/+^* and *Chmp5^Ctsk^;Rosa26^mTmG/+^* mice, respectively. *n* = 3 with tissues from 3 different animals per group. Scale bars, 100 μm; arrows indicating the periosteum. (**F**) Gross image demonstrating aging-related phenotypes in *Chmp5^Ctsk^* compared to *Chmp5^Ctsk/+^* mice at 1 year of age. Images are representative of 5 animals per group. (**G**) Cell cycle analysis in ATDC5 cells with or without *Chmp5* deletion. *n* = 3 replicates per group, results repeated twice. All data are mean ± s.d.; 2-way ANOVA followed by multiple comparisons or two-tailed unpaired Student’s *t*-test (E).

Notably, the low apoptotic cell ratio (approximately 5%) could not well match the decrease in cell proliferation rate (approximately 58%) in *Chmp5^Ctsk^* periskeletal progenitors. We further analyzed the RNA-seq data of *Chmp5^Ctsk^* vs. *Chmp5^Ctsk/+^* cells to dig out the mechanism of CHMP5 deficiency on the fate of skeletal progenitors. The gene set enrichment analysis (GSEA) showed significant enrichment of genes in multiple Reactome molecular pathways associated with cell senescence in *Chmp5^Ctsk^* relative to *Chmp5^Ctsk/+^* periskeletal progenitors, including *Hmga1*, *Hmga2*, *Trp53*, *Ets1*, and *Txn1* (**Fig. 3B and C**). Meanwhile, GSEA of RNA-seq data also showed significant enrichment of the SAUL_SEN_MAYO geneset (positively correlated with cell senescence) and the KAMMINGA_SENESCENCE geneset (negatively correlated with cellular senescence) in *Chmp5^Ctsk^* vs. *Chmp5^Ctsk/+^* periskeletal progenitors (**Fig. S3C**).

Furthermore, Western blot results showed that both the p16 and p21 proteins, two canonical markers of cellular senescence, were upregulated in *Chmp5^Ctsk^* compared to control cells, with a higher degree of upregulation in the p16 protein (**Fig. 3D**). However, the mRNA levels of *Cdkn2a* (p16) and *Cdkn1a* (p21) did not show significant changes according to the RNA-seq analysis (**Fig. S3D**). Furthermore, immunostaining for another cell senescence marker γH2Ax demonstrated that there were significantly more γH2Ax^+^;GFP^+^ cells in periskeletal overgrowth in *Chmp5^Ctsk^*;*Rosa26^mTmG/+^* mice relative to the periosteum of *Chmp5^Ctsk/+^*;*Rosa26^mTmG/+^* control mice (**Fig. 3E**). Accordingly, C*hmp5^Ctsk^* mice showed gross accelerated aging-related phenotypes, including hair loss, joint stiffness/contracture, and decreased motility (**Fig. 3F**).

To further characterize the effect of CHMP5 on skeletal progenitor cell fate, we deleted *Chmp5* in ATDC5 cells, which were shown to be an appropriate model of periskeletal progenitors in a previous study [23], using CRISPR/CAS9 technology (**Fig. S3E**). Similarly, *Chmp5* deletion also significantly suppressed the cell proliferation rate of ATDC5 cells and mildly increased cell apoptosis (**Fig. S3F and G**). However, inhibition of cell apoptosis using the pancaspase inhibitor Q-VD-Oph could not reverse cell number reduction (**Fig. S3H**), indicating that cell apoptosis was not the main cause of the decrease in the proliferation rate of *Chmp5*-deficient skeletal progenitors. Instead, cell cycle analysis showed a significant decrease of *Chmp5*-deficient ATDC5 cells in phase S, and a significant proportion of these cells were arrested in G0/G1 or G2/M phases (**Fig. 3G**), which is a typic characteristic of cell cycle arrest in senescent cells. Taken together, these results at molecular, cellular, and gross levels demonstrate that *Chmp5* deficiency induces skeletal progenitor cell senescence.

### Secretory phenotype of *Chmp5*-deficient skeletal progenitors

A remarkable feature of senescent cells is the senescence associated secretory phenotype (SASP), which plays a critical role in mediating the pathophysiological processes of cell senescence [24]. In particular, periskeletal overgrowths in *Chmp5^Ctsk^*;*Rosa26^mTmG/+^* mice showed evident integration of many cells lacking evidence of Cre-mediated recombination (GFP^−^) (**Fig. 1E**), suggesting that *Chmp5*-deleted periskeletal progenitors might recruit neighboring *wild-type* cells to facilitate the bone overgrowth. Additionally, the GSEA of RNA-seq data showed a significant enrichment of genes associated with the SASP molecular pathway in *Chmp5*-deleted periskeletal progenitors (**Fig. 4A**). Meanwhile, confocal fluorescence microscopy captured many extracellular vesicles shed from the plasma membrane of *Chmp5^Ctsk^* periskeletal progenitors, which were rarely detected around *wild-type* control cells (**Fig. 4B**). Furthermore, nanoparticle tracking analysis demonstrated higher concentrations of extracellular vesicles in the culture medium of *Chmp5^Ctsk^* periskeletal progenitors compared to *wild-type* control cells (**Fig. 4C, D** and **Fig. S4A**). These results show that *Chmp5* deficiency activates the SASP molecular pathway and increases the secretion of skeletal progenitors.

**Fig. 4.**
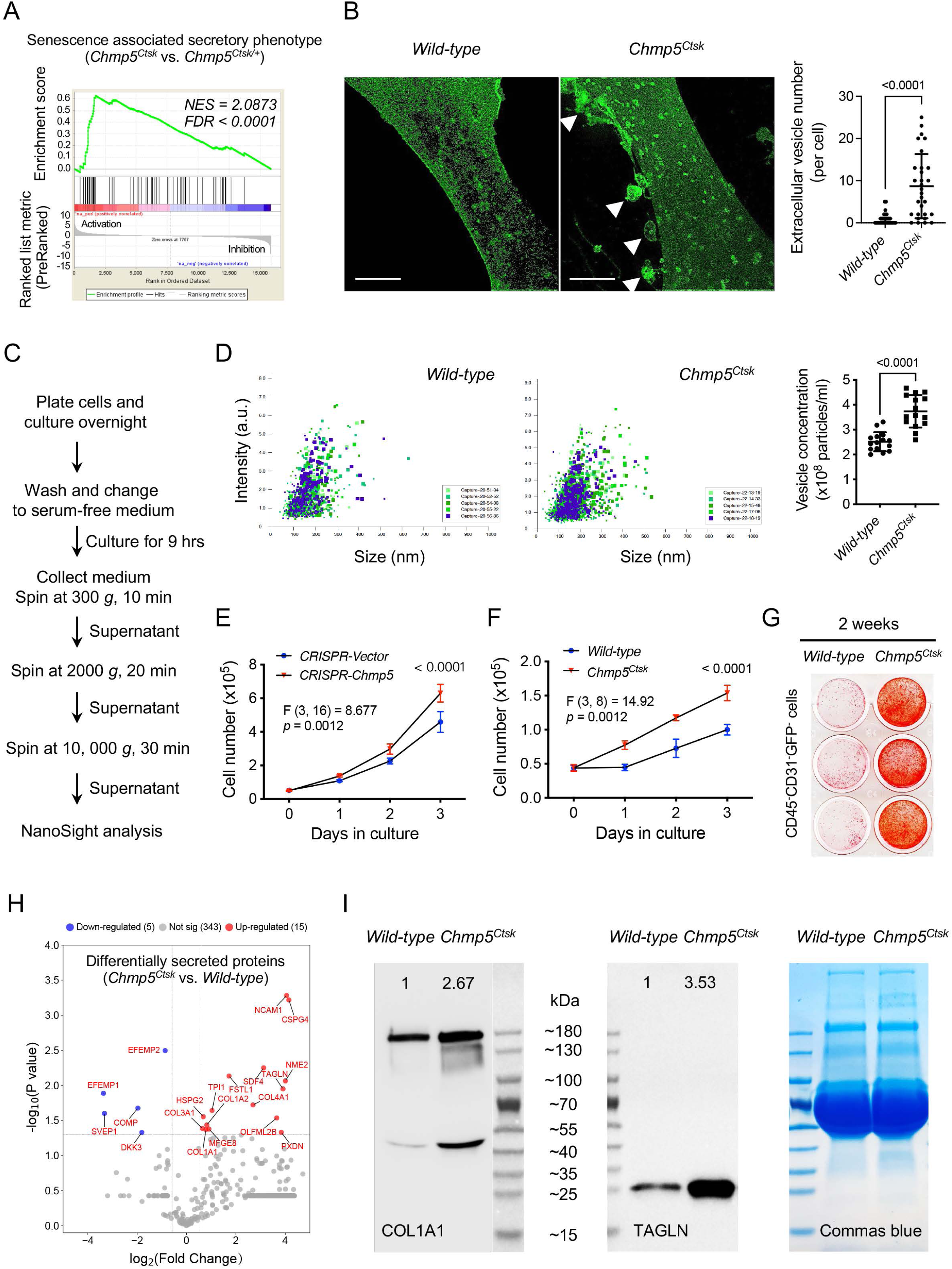
Secretory phenotype of *Chmp5*-deficient skeletal progenitors. (**A**) GSEA of RNA-seq data showing positive enrichment of genes associated with the molecular pathway of senescence-associated secretory phenotype (Reactome database) in *Chmp5^Ctsk^* relative to *Chmp5^Ctsk/+^* periskeletal progenitors. (**B**) Confocal images showing and quantifying extracellular vesicles (arrowheads) around *Chmp5^Ctsk^* periskeletal progenitors. Images are representative of 30 cells from 3 animals per group. Scale bars, 10 μm. (**C** and **D**) Nanoparticle tracking analysis of extracellular vesicles in culture medium of *Chmp5^Ctsk^* and *wild-type* periskeletal progenitors. Data pooled from 3 replicates per group, 5 reads for each; repeated twice using cells from 2 mice per group. (**E**) Cell number counting determining cell proliferation in ATDC5 cells treated with culture supernatant from *Chmp5*-deficient or *Chmp5*-sufficient cells. *n* = 3 per group per time point, repeated twice. (**F** and **G**) Cell number counting and alizarin red staining examining cell proliferation (**F**) and osteogenesis (**G**) in neighboring CD45^−^;CD31^−^;GFP-progenitors sorted from periskeletal tissues of *Chmp5^Ctsk^;Rosa26^mTmG/+^* or *Ctsk-Cre;Rosa26^mTmG/+^* mice. *n* = 3 replicates per group per time point, repeated 3 times using cells from 3 different animals. (**H**) Volcano plot showing differentially secreted protein in supernatants of *Chmp5^Ctsk^* vs. *wild-type* periskeletal progenitors analyzed by Nano LC-MS/MS. *n* = 3 with cells from 3 different animals per group. (**I**) Western blot verifying the increase in COL1A1 and TAGLN proteins in supernatants of *Chmp5^Ctsk^* relative to *wild-type* periskeletal progenitors. Commas blue stain as the loading control. Results repeated 3 times using cells from 3 different mice each group. All data are mean ± s.d.; two-tailed unpaired Student’s *t*-test for comparison of two groups or 2-way ANOVA followed by multiple comparisons.

Functionally, conditioned medium collected from *Chmp5*-deficient skeletal progenitors caused a higher proliferation rate of ATDC5 cells than medium from control cells (**Fig. 4E**). Meanwhile, coculture of *wild-type* with *Chmp5^Ctsk^* periskeletal progenitors promoted osteogenesis of *wild-type* cells (**Fig. S4B**). Simultaneously, CD45^−^;CD31^−^;GFP-skeletal progenitors from periskeletal tissues of *Chmp5^Ctsk^*;*Rosa26^mTmG/+^* mice showed increased proliferation along with enhanced osteogenic differentiation compared to the corresponding cells from *Ctsk-Cre;Rosa26^mTmG/+^* control mice (**Fig. 4F, G** and **Fig. S4C, D**). Immunostaining for the cell proliferation marker Ki-67 demonstrated an increase in the number of stained cells in the periskeletal lesion of *Chmp5^Ctsk^* mice (**Fig. S4E**). Since GFP^+^ *Chmp5^Ctsk^* periskeletal progenitors showed a decreased cell proliferation rate and increased cell senescence (**Fig. 3**), and the GFP-*Chmp5^Ctsk^* periskeletal progenitors showed an increased cell proliferation rate (**Fig. 4F** and **Fig. S4C**), these Ki-67^+^ cells in **Fig. S4E** should represent GFP- skeletal progenitors in the periskeletal lesion. Together, these results demonstrate that *Chmp5*-deficient skeletal progenitors could promote the proliferation and osteogenesis of surrounding *wild-type* skeletal progenitors.

Next, we performed a secretome analysis to characterize secretory profiles of *Chmp5^Ctsk^* vs. *wild-type* periskeletal progenitors. Fifteen proteins were found to increase and five proteins to decrease in the cell supernatant of *Chmp5^Ctsk^* periskeletal progenitors (**Fig. 4H** and **Fig. S4F**). In particular, all 15 upregulated proteins, including NCAM1, CSPG4, SDF4, NME2, TAGLN, OLFML2B, PXDN, COL4A1, FSTL1, TPI1, MFGE8, COL1A2, COL1A1, HSPG2, and COL3A1, have been identified in the osteoblastic cell secretome or in the regulation of osteoblast differentiation or functions [25, 26]. Western blot verified upregulation of two of these proteins, COL1A1 and TAGLN, in the cell supernatant of *Chmp5^Ctsk^* vs. *wild-type* periskeletal progenitors (**Fig. 4I**). Notably, the secretome analysis did not detect common SASP factors, such as cytokines and chemokines, in the secretory profile of *Chmp5^Ctsk^* periskeletal progenitors, probably due to their small molecular weights and the technical limitations of the mass-spec analysis. Taken together, these results demonstrate a secretory phenotype and paracrine actions of *Chmp5*-deficient skeletal progenitors. However, factors that mediate the paracrine actions of *Chmp5*-deficient periskeletal progenitors remain further clarified in future studies.

### Senolytic treatment mitigates musculoskeletal pathologies in *Chmp5* conditional knockout mice

To affirm the role of cell senescence of osteogenic cells in contributing to bone overgrowth in *Chmp5^Ctsk^* and *Chmp5^Dmp1^* mice, we treated *Chmp5*-dificient periskeletal progenitors and animals with the senolytic drugs quercetin and dasatinib (Q + D), which have been widely used to eliminate senescent cells under various physiological and pathological conditions [27, 28], and assessed the therapeutic efficacy. First, in vitro Q + D treatment showed that *Chmp5^Ctsk^* periskeletal progenitors were more sensitive to senolytic drugs for cell apoptosis compared to control cells (**Fig. 5A**). *Chmp5^Ctsk^* animals were then treated with Q + D from the first two weeks after birth for 7 weeks and showed a significantly improved periskeletal bone overgrowth and animal motility compared to *Chmp5^fl/fl^* control animals (**Fig. 5B** and **Fig. S5**). Similarly, treatment of *Chmp5^Dmp1^* mice with Q + D also improved musculoskeletal manifestations, including hindlimb abduction, the whole femur and cortical bone thickness, and partially restored skeletal muscle functions (**Fig. 5C-E**). An improvement in animal motility was also found with Q + D treatment in adult mice. There were no apparent abnormalities in *Chmp5^fl/fl^* control animals treated with the same dose of Q + D in parallel.

**Fig. 5.**
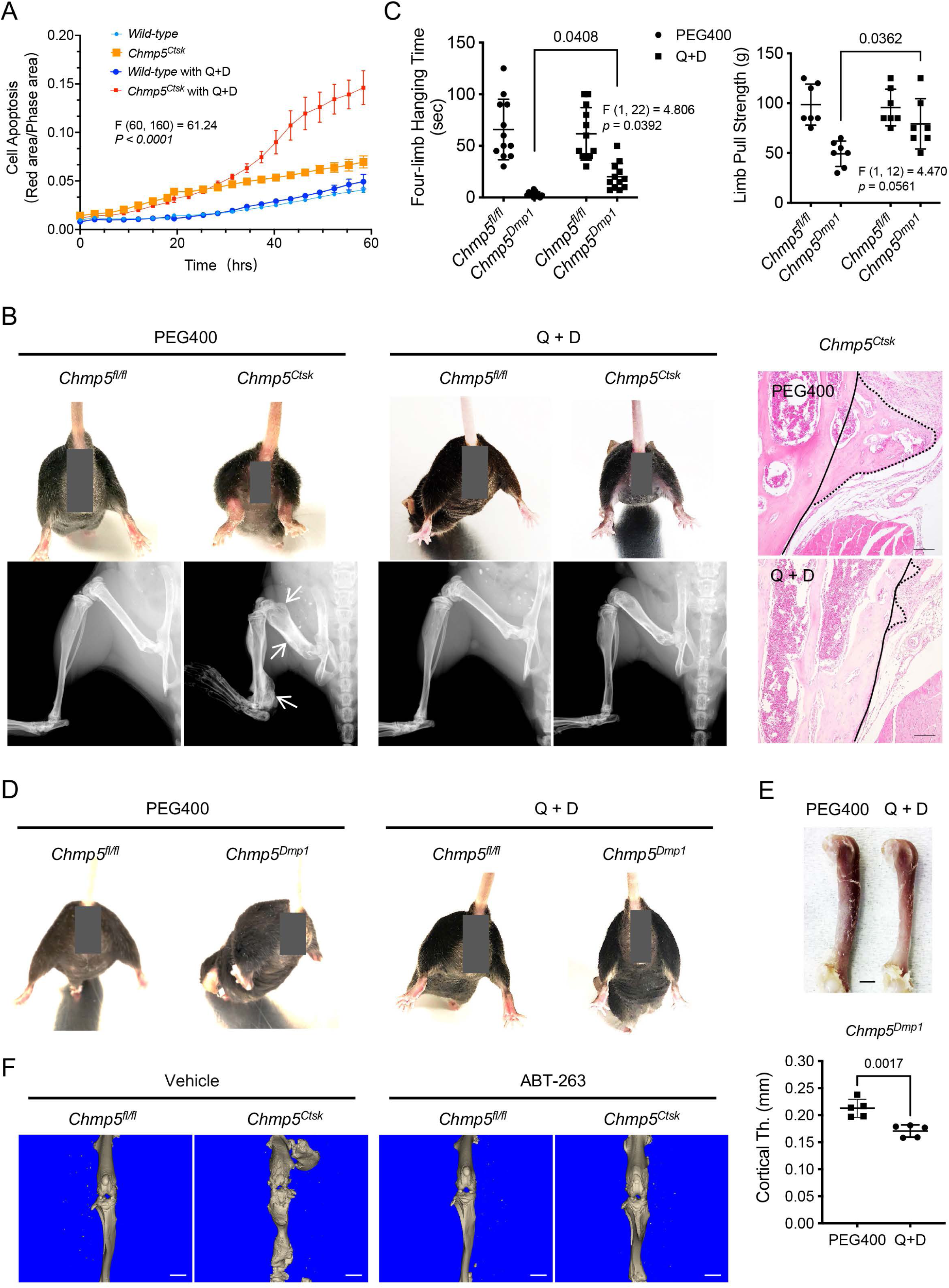
Senolytic treatment mitigates musculoskeletal pathologies in *Chmp5* conditional knockout mice. (**A**) Incucyte live-cell apoptosis analysis of *Chmp5^Ctsk^* and *wild-type* periskeletal progenitors treated with 50 μM Quercetin and 500 nM dasatinib (Q + D) and labeled by Annexin V. *n* = 6 for each group, experiment repeated twice using cells from different animals. (**B**) Gross images, radiography, and histology (H&E) respectively demonstrating hindlimb abduction and periskeletal bone overgrowth (arrows or dot-line) in *Chmp5^Ctsk^* in comparison with *Chmp5^fl/fl^* mice after treatment with Q + D or vehicle PEG400 weekly for 7 weeks. *n* = 8-10 mice per group. Continuous lines in H&E images indicating an approximate edge of the cortical bone and dot lines indicating the edge of periskeletal overgrowing bones. Scale bars, 100 μm. (**C**) Four-limb hanging time and forelimb pull strength in *Chmp5^Dmp1^* and *Chmp5^fl/fl^* mice after treatment with Q + D or the vehicle PEG400 weekly for 16 weeks. *n* = 12 animals per group for hanging time test pooled from both genders; *n* = 7 male mice per group for the forelimb pull strength test, similar changes found in both genders. (**D**) Gross images showing hindlimb abduction in *Chmp5^Dmp1^* compared to *Chmp5^fl/fl^* mice after treatment with Q + D or the vehicle PEG400. *n* = 12 animals per group. (**E**) Gross image and Micro-CT analysis demonstrating femur and cortical bone thickness in *Chmp5^Dmp1^* mice after treatment with Q + D or the vehicle PEG400. *n* = 5 male mice per group, similar changes found in both genders. Scale bar, 1 mm. (**F**) Micro-CT images showing periskeletal bone overgrowth in *Chmp5^Ctsk^* in comparison with *Chmp5^fl/fl^* mice after treatment with ABT-263 or vehicle weekly for 8 weeks. *n* = 4 mice per group, Scale bars, 2 mm. All data are mean ± s.d.; 2-way ANOVA followed by multiple comparisons or two-tailed unpaired Student’s *t*-test for comparison of two groups.

Furthermore, to verify the efficacy of senolytic drugs in treating the periskeletal bone overgrowth in *Chmp5^Ctsk^* mice, we used another senolytic drug Navitoclax (ABT-263), which is a BCL-2 family inhibitor and specifically induces apoptosis of senescent cells [29, 30]. Micro-CT analysis demonstrated that ABT-263 could also relieve periskeletal bone overgrowth in *Chmp5^Ctsk^* mice (**Fig. 5F**). Together, these results demonstrate that the elimination of senescent cells using senolytic drugs is effective in alleviating musculoskeletal pathologies in *Chmp5^Ctsk^* and *Chmp5^Dmp1^* animals, confirming that osteogenic cell senescence is responsible for musculoskeletal abnormalities in *Chmp5^Ctsk^* and *Chmp5^Dmp1^* mice.

### CHMP5 is essential for endolysosomal functions and maintaining VPS4A protein in skeletal progenitors

Next, we wondered whether *Chmp5* deficiency in skeletal progenitors affects the endocytic pathway. Notably, approximately 45% of *Chmp5^Ctsk^* periskeletal progenitors contained enlarged GFP^+^ vesicles, which were rarely found in *wild-type Ctsk^+^* periskeletal progenitors (**Fig. 6A**), suggesting a possible issue of the endocytic pathway in *Chmp5*-deleted skeletal progenitors. Next, we used molecular markers for different components and the function of the endocytic pathway to elucidate the influence of *Chmp5* deficiency on the endolysosomal system in skeletal progenitors (**Fig. 6B**).

**Fig. 6.**
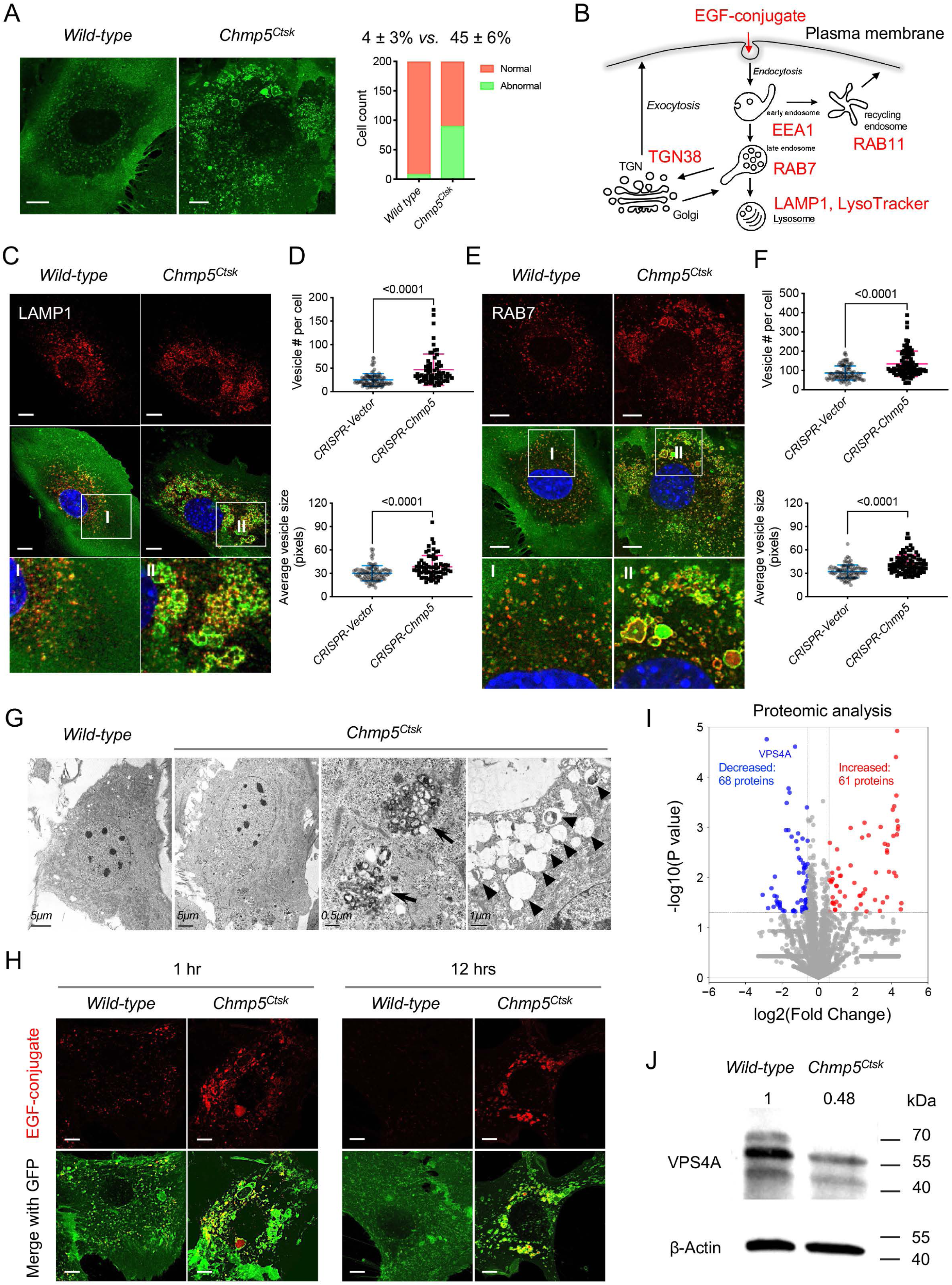
CHMP5 is essential for endolysosomal functions and maintaining VPS4A protein in skeletal progenitors. (**A**) Representative confocal images demonstrating abnormally enlarged vesicles in cultured *Chmp5^Ctsk^* relative to *wild-type* periskeletal progenitors. Abnormal cells identified by containing enlarged GFP^+^ vesicles. *n* = 200 cells from 3 mice per group. (**B**) Schematic showing molecular markers utilized to analyze the endocytic pathway. (**C**) Representative confocal images showing LAMP1 immunostaining in *Chmp5^Ctsk^* and *wild-type* periskeletal progenitors. *n* = 20 cells per group. (**D**) Quantification of LAMP1^+^ vesicles in ATDC5 cells with or without *Chmp5* depletion. *n* = 93, 66 cells, respectively. (**E**) Representative confocal images showing RAB7 immunostaining in *Chmp5^Ctsk^* and *wild-type* periskeletal progenitors. *n* = 15 cells per genotype. (**F**) Quantification of RAB7^+^ vesicles in ATDC5 cells with or without *Chmp5* depletion. *n* = 94, 92 cells, respectively. (**G)** Transmission electron microscopy showing the accumulation of multivesicular body-like structures (arrows) and lysosome-like structures (arrowheads) in *Chmp5^Ctsk^* relative to *wild-type* periskeletal progenitors. *n* = 30 cells per group. (**H**) Confocal live cell images demonstrating delayed degradation of the EGF conjugate in *Chmp5^Ctsk^* vs. *wild-type* periskeletal progenitors. *n* = 10 cells each group per time point. (**I**) Volcano plot of proteomic analysis showing differentially expressed proteins in *Chmp5^Ctsk^* vs. *wild-type* periskeletal progenitors. *n* = 3 with cells from 3 different animals. (**J**) Western blot verifying the decrease in VPS4A protein in *Chmp5^Ctsk^* vs. *wild-type* periskeletal progenitors. The numbers above lanes indicating the intensity of the VPS4A band relative to that of *wild-type* cells after normalization by β-Actin. Results repeated 3 times with cells from 3 different mice each group. All data are mean ± s.d.; Mann-Whitney test for the comparison of the vesicle number and two-tailed unpaired Student’s *t*-test for the comparison of the vesicle size in (**D**) and (**F**). Scale bars, 10 μm except (**G**) as indicated.

The LysoTracker Red DND-99 trace and the immunostaining of lysosome-associated membrane protein 1 (LAMP1) showed positive staining in many of the enlarged vesicles in *Chmp5^Ctsk^* periskeletal progenitors (**Fig. 6C** and **Fig. S6A, B**). The quantification of the number and average size of LAMP1 stained vesicles were further performed in ATDC5 cells with or without *Chmp5* deletion (**Fig. 6D**). Next, the number and size of the vesicles stained for the late endosome marker RAB7 also increased markedly in *Chmp5*-deficient relative to *wild-type* skeletal progenitors (**Fig. 6E, F**). Meanwhile, many of these accumulated vesicles were double positive for LAMP1 and RAB7 (**Fig. S6C, D**), suggesting that they are terminal compartments in the endocytic pathway. Accordingly, the transmission electron microscopy demonstrated a significant accumulation of MVB-like and lysosome-like structures with different electronic density in *Chmp5^Ctsk^* periskeletal progenitors (**Fig. 6G**).

To examine the function of the endocytic pathway, we used a pH-sensitive fluorescent EGF conjugate and traced the degradation of the internalized EGF receptor in *Chmp5^Ctsk^* vs. *wild-type* periskeletal progenitors. *Chmp5* deficiency significantly delayed the degradation of the EGF conjugate in *Chmp5^Ctsk^* periskeletal progenitors (**Fig. 6H**), indicating altered function of the endocytic pathway. It should be noted that the early endosomes of EEA1^+^ also increased slightly in approximately 10% of *Chmp5^Ctsk^* periskeletal progenitors, while the recycling endosomes of RAB11^+^ and the TGN38^+^ trans-Golgi network were relatively normal (**Fig. S6E**). These results demonstrate that CHMP5 deficiency in skeletal progenitors causes the accumulation of late endosomes and lysosomes and disrupts the function of the endolysosomal pathway.

To dissect the molecular mechanism of CHMP5 that regulates the endolysosomal pathway, we performed a proteomic analysis in *Chmp5^Ctsk^* and *wild-type* periskeletal progenitors. Bioinformatic analyses of proteomic data showed that VPS4A is one of the most significant decreased proteins in *Chmp5^Ctsk^* compared to *wild-type* periskeletal progenitors (**Fig. 6I**). The decrease in the VPS4A protein in *Chmp5^Ctsk^* periskeletal progenitors was verified by Western blot analysis (**Fig. 6J**), while the *Vps4a* mRNA did not show a significant change in the RNA-seq analysis (**Fig. S6F**). On the other hand, when CHMP5 was overexpressed by transfection of a CHMP5 vector into HEK-293T cells, the level of VPS4A protein increased significantly and *VPS4A* mRNA was not significantly affected (**Fig. S6G, H**). These results indicate that CHMP5 is essential in maintaining the VPS4A protein. Since VPS4A is a critical ATPase mediating the function of the ESCRT-III protein complex, the reduction of VPS4A protein could be responsible for endolysosomal dysfunction in *Chmp5*-deleted skeletal progenitors.

### Mitochondrial dysfunction is responsible for cell senescence in *Chmp5*-deleted skeletal progenitors

A remaining question is how *Chmp5* deficiency and endolysosomal dysfunction cause skeletal cell senescence. Impairment of lysosomal functions could hinder the degradation of damaged mitochondria, resulting in the accumulation of reactive oxygen species (ROS) in mitochondria [31, 32]. Indeed, the GSEA of RNA-seq data showed significant enrichment of genes associated with the molecular pathway of oxidative stress-induced cell senescence in *Chmp5^Ctsk^* vs. *Chmp5^Ctsk/+^* periskeletal progenitors (**Fig. 7A**). Accordingly, the level of mitochondrial ROS and the abundance of mitochondria increased markedly in *Chmp5*-deficient compared to *wild-type* skeletal progenitors, as shown by CellRox, tetramethylrhodamine methyl ester (TMRE), and MitoTracker fluorescence stain, and expression levels of mitochondrial inner membrane proteins NDUF88, SDHB, and ATP5A (**Fig. 7B to D and Fig. S7A, B**).

**Fig. 7.**
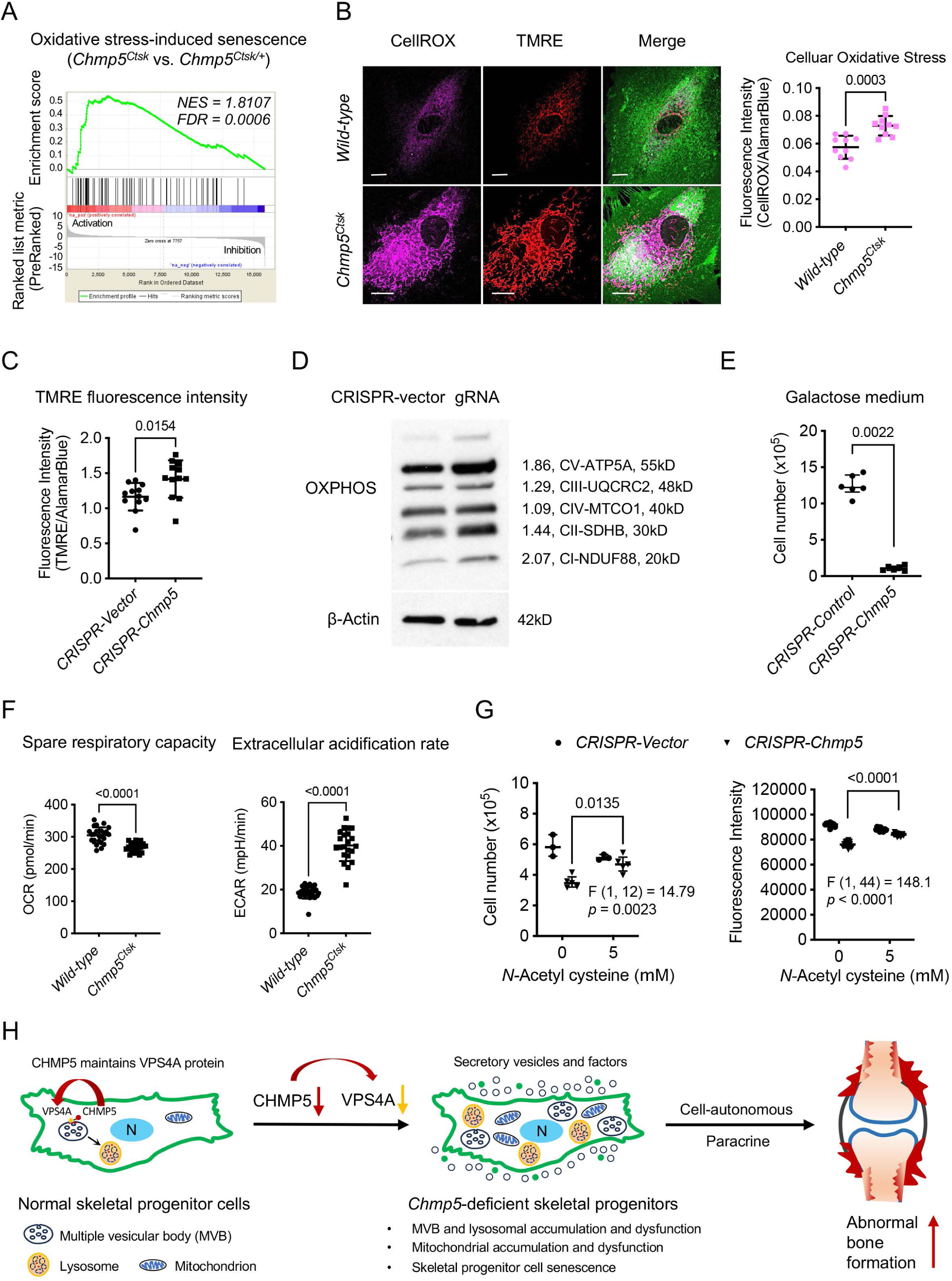
Mitochondrial dysfunction is responsible for cell senescence in *Chmp5*-deleted skeletal progenitors. (**A**) GSEA of RNA--seq data reporting positive enrichment of genes associated with the molecular pathway of oxidative stress-induced senescence in *Chmp5^Ctsk^* relative to *Chmp5^Ctsk/+^* periskeletal progenitors. (**B**) Confocal fluorescence images showing intracellular ROS (CellROX Deep Red) and mitochondria (TMRE) in *Chmp5^Ctsk^* vs. *wild-type* periskeletal progenitors. Images are representative of 30 cells per group. Scale bars, 15 μm. The graph on the right shows the quantification of the fluorescence intensity of CellROX Deep Red. *n* = 10 each group, results repeated twice. (**C**) Quantification of TMRE fluorescence intensity in *Chmp5*-sufficient and *Chmp5*-deficient ATDC5 cells. *n* = 12 for each group, repeated 3 times. (**D**) Western blotting determining the expression of mitochondrial OXPHOS proteins NDUF88, SDHB, MT-CO1, UQCRC2, and ATF5A in *Chmp5*-deficient compared to *Chmp5*-sufficient ATDC5 cells. Experiment repeated twice. (**E**) Cell number counting to determine cell proliferation in galactose medium. *n* = 6 replicates per group, repeated three times. (**F**) Seahorse mitochondrial stress test showing mitochondrial respiratory capacity and extracellular acidification rate in *Chmp5^Ctsk^* relative to *wild-type* periskeletal progenitors. Data pooled from cells of 3 mice, 8 replicates for each sample. (**G**) Cell number counting and AlamarBlue assay determining cell proliferation in ATDC5 cells with or without *Chmp5* depletion after treatment with the antioxidant *N*-Acetyl cysteine. *n* = 3 or 12 each group for cell number counting or AlamarBlue assay, respectively; results repeated 3 times. (**H**) Schematic showing the function of CHMP5 in maintaining VPS4A protein and endolysosomal homeostasis and restricting cell senescence and osteogenesis in skeletal progenitors. All data are mean ± s.d.; two-tailed unpaired Student’s *t*-test, except the Mann-Whitney test in panel (**E**).

Mitochondrial functions were also compromised in *Chmp5*-deficient skeletal progenitors as shown by decreased mitochondrial respiratory capacity, increased extracellular acidification rate, and reduced tolerance to galactose (**Fig. 7E and F**). Furthermore, mitochondrial dysfunction and DNA damage response usually form a feedback loop to drive senescent cell phenotypes [33–35]. The GSEA of the RNA-seq data also showed enrichment of genes associated with DNA damage-induced cell senescence in *Chmp5^Ctsk^* vs. *Chmp5^Ctsk/+^* skeletal progenitors (**Fig. S7C**). Upregulation of two critical DNA damage-responsive genes *H2afx* and *Trp53* was verified by qPCR in *Chmp5^Ctsk^* relative to *Chmp5^Ctsk/+^* skeletal progenitors (**Fig. S7D**). Importantly, N-Acetylcysteine (NAC) antioxidant treatment could reverse the reduced activity of cell proliferation in *Chmp5*-deficient skeletal progenitors (**Fig. 7G**). Therefore, the accumulation of dysfunctional mitochondria and mitochondrial ROS is responsible for cell senescence in *Chmp5*-deficient skeletal progenitors.

## Discussion

In this study, our results reveal the role and mechanism of CHMP5 in the regulation of cell senescence and bone formation in osteogenic cells. In normal skeletal progenitor cells, CHMP5 is essential in maintaining the VPS4A protein, a critical AAA ATPase for the functions of the ESCRT-III protein complex. The deficiency of CHMP5 decreases the VSP4A protein and causes the accumulation of dysfunctional late endosomes (MVB), lysosomes, and mitochondria, which induce skeletal cell senescence and result in increased bone formation through a combination of cell-autonomous and paracrine mechanisms (**Fig. 7H**). Strikingly, elimination of senescent cells using senolytic drugs is effective in mitigating abnormal bone formation.

The role of CHMP5 in cell senescence has not been reported. As the mechanisms and functions of cell senescence could be highly heterogeneous depending on inducers, tissue and cell contexts, and other factors such as “time” [36–38], there is currently no universal molecular marker to define all types of cell senescence. In this study, *Chmp5*-deficient skeletal progenitors show multiple canonical features of senescent cells, including increased p16 and p21 proteins and γH2Ax^+^ cells, cell cycle arrest, elevated secretory phenotype, accumulation and dysfunction of lysosomes and mitochondria. Also, *Chmp5^Ctsk^* and *Chmp5^Dmp1^* mice show dramatic aging-related phenotypes, including hair loss, joint stiffness/contracture, decreased bone strength, or gradual loss of muscle mass and functions. Importantly, senolytic drugs markedly improve musculoskeletal abnormalities and increase animal motility in *Chmp5^Ctsk^* and *Chmp5^Dmp1^* mice. These results reveal the role of CHMP5 in controlling the senescence of skeletal cells.

CHMP5 has been reported to regulate VPS4 activity, which catalyzes the disassembly of the ESCRT-III protein complex from late endosomes/MVBs, by binding directly to the VPS4 protein or other protein partners such as LIP5 and Brox [39, 40]. However, it remains unknown how CHMP5 regulates VPS4 activity. Our results in this study show that CHMP5 is necessary to maintain the VPS4A protein but does not affect the level of VPS4A mRNA, suggesting that CHMP5 regulates VPS4 activity by maintaining the level of the VPS4A protein. However, the mechanism of CHMP5 in the regulation of the VPS4A protein has not yet been studied. Since CHMP5 can recruit the deubiquitinating enzyme USP15 to stabilize IκBα in osteoclasts by suppressing ubiquitination-mediated proteasomal degradation [8], it is also possible that CHMP5 stabilizes the VPS4A protein by recruiting deubiquitinating enzymes and regulating the ubiquitination of VPS4A, which needs to be clarified in future studies. Notably, mutations in the *VPS4A* gene in humans can cause multisystemic diseases, including musculoskeletal abnormalities [41] (OMIM: 619273), suggesting that normal expression and function of VPS4A are important for musculoskeletal physiology. The roles of VPS4A in regulating musculoskeletal biology and cell senescence should be further explored in future studies.

The upstream signaling that regulates CHMP5 expression in osteogenic cells is unclear. Previous studies showed that muramyl-dipeptide and lipopolysaccharide upregulate CHMP5 expression in human monocytes [42, 43]. Whether the mechanism also occurs in osteogenic cells remains to be determined, especially since many molecular and cellular mechanisms are cell lineage/type dependent. In addition, an earlier study showed that *Rank* haploinsufficiency in *Chmp5^Ctsk^* mice (*Chmp5^Ctsk^;Rank^+/−^*) could greatly reverse skeletal abnormalities in these animals, including periskeletal overgrowth [8]. Therefore, RANK receptor signaling may function upstream of CHMP5. However, whether RANK is expressed in osteogenic cells needs additional studies to clarify. Otherwise, other epigenetic, transcriptional, translational, or post-translational mechanisms may also play a role in the regulation of CHMP5 expression in osteogenic lineage cells in physiological and pathological contexts.

A previous study using the *Chmp5^Ctsk^* mouse model defined a function of CHMP5 in the regulation of osteoclast differentiation by tuning NF-κB signaling [8]. While the evidence on the role of CHMP5 in osteoclastogenesis is robust in that study, our results in the current study provide compelling evidence on the role of CHMP5 in bone formation in osteogenic cells. Furthermore, the function of CHMP5 in restricting bone formation was confirmed in the *Chmp5^Dmp1^* mouse model and in the MC3T3-E1 cell model (Figs. 1 and 2). Therefore, it is probable that both osteoclast and osteoblast lineage cells contribute to bone deformation in *Chmp5^Ctsk^* mice. However, the previous study by Greenblatt et al. did not find endolysosomal abnormalities in osteoclasts after *Chmp5* deletion, and in that study cell senescence and mitochondrial functions were not measured. Also, a previous study by Adoro et al. did not detect endolysosomal abnormalities in *Chmp5*-deficient developmental T cells [9]. Since both osteoclasts and T cells are of hematopoietic origin, and meanwhile osteogenic cells and MEFs, which show endolysosomal abnormalities after CHMP5 deficiency, are of mesenchymal origin, it turns out that the function of CHMP5 in regulating the endolysosomal pathway could be cell lineage-specific, which remains clarified in future studies.

Furthermore, it is unclear whether the effect of senolytic drugs in *Chmp5^Ctsk^* mice involves targeting osteoclasts other than osteogenic cells, as osteoclast senescence has not yet been evaluated. However, the efficacy of Q + D in targeting osteogenic cells, which is the focus of the current study, was confirmed in *Chmp5^Dmp1^* mice (Fig. 5C-E). Additionally, Q + D caused a higher cell apoptotic ratio in *Chmp5^Ctsk^* compared to *wild-type* periskeletal progenitors in *ex vivo* culture (Fig. 5A), demonstrating the effectiveness of Q + D in targeting osteogenic cells in the *Chmp5^Ctsk^* model. Furthermore, an alternative senolytic drug ABT-263 could also ameliorate periskeletal bone overgrowth in *Chmp5^Ctsk^* mice (Fig. 5F). Together, these results confirm that osteogenic cell senescence is responsible for the bone overgrowth in *Chmp5^Ctsk^* and *Chmp5^Dmp1^* mice and senolytic treatments are effective in alleviating these skeletal disorders.

Notably, aging is associated with decreased osteogenic capacity in marrow stromal cells, which is related to conditions with low bone mass, such as osteoporosis. Rather, aging is also accompanied by increased ossification or mineralization in musculoskeletal soft tissues, such as tendons and ligaments [44]. In particular, the abnormal periskeletal overgrowth in *Chmp5^Ctsk^* mice was predominantly mapped to the insertion sites of tendons and ligaments on the bone (Fig. 1A and E), which is consistent with changes during aging and suggests that mechanical stress at these sites could contribute to the aberrant bone growth. These results suggest that skeletal stem/progenitor cells at different sites of musculoskeletal tissues could demonstrate different, even opposite outcomes in osteogenesis, due to cell senescence.

In addition, there was also a mild increase in apoptotic cells in the population of *Chmp5*-deficient skeletal progenitors (approximately 5%). Since anti-apoptotic treatment with the pan-caspase inhibitor Q-VD-Oph could not reverse the decrease in cell number in *Chmp5*-deficient skeletal progenitors, cell apoptosis may not be the main cause of the decrease in cell proliferation rate in these progenitors. It should be noted that although senescent cells are generally resistant to apoptosis, the gradual accumulation of dysfunctional lysosomes and mitochondria in *Chmp5*-deficient skeletal progenitors could result in overload of toxic macromolecules and metabolites, which could activate the apoptotic pathway to eliminate severely damaged cells. Otherwise, CHMP5 may regulate cell senescence and apoptosis through different mechanisms.

In summary, we have revealed the role of CHMP5 in the regulation of cell senescence and bone formation in osteogenic cells. This study also confirms the essential function of CHMP5 in the endolysosomal pathway that involves the regulation of the VPS4A protein. Additionally, mitochondrial functions are impaired with CHMP5 deletion, which is responsible for cell senescence. Future studies should explore the mechanism of CHMP5 in the regulation of mitochondrial functions. Since *Chmp5^Ctsk^* and *Chmp5^Dmp1^* mice recapitulate many cellular and phenotypic features of musculoskeletal lesions in lysosomal storage disease, these animals could be used as preclinical models to investigate the mechanism and test therapeutic drugs for these intractable disorders.

## Acknowledgments

We thank Dr. Sankar Ghosh for providing *Chmp5*-floxed mice; Drs. Gregory Hendricks and Lara Strittmatter at the University of Massachusetts Chan Medical School for help in interpreting TEM data; Dr. Tomer Shpilka for assistance in running the Seahorse analyzer; Drs. Guangchuang Yu and Jiyeon Park for independently analyzing the RNA-seq data. Dr. Chunjing Bian for generating the CHMP5 overexpression vector. X.G. is supported by the National Natural Science Foundation of China (project 2-2-008-0213) and the Beijing Natural Science Foundation (project 5222008). J.H.S. holds support from the NIH (R21AR077557, R01AR078230) and AAVAA Therapeutics.

## Contributions

X.G. perceived the project, designed experiments, interpreted the results, and wrote and reviewed the manuscript; F.Z., Y.W., L.Z., C.W. and X.G. performed experiments and analyzed the results; D.C. and R.X. provided critical resources for the study; H.L. contributed to part of RNA-seq data analyses; C.M.H. offered suggestions and main reagents for mitochondrial analyses; J.H.S. partly supported the study and reviewed the manuscript.

## Methods

### Mice

In this study, two *Chmp5*-flox mouse strains were used, and similar phenotypes were observed. The *Chmp5^tm2.1Gho^* strain was previously described [8]. The alternative *Chmp5*-flox strain was established by CRISPR/Cas9-mediated genome engineering. Similarly to the *Chmp5^tm2.1Gho^* strain, exons 4 and 5 were selected as the conditional knockout region in the newly created strain and deletion of this region results in frameshift of the *Chmp5* gene. To engineer the target vector, homologous arms, and the conditional knockout region with 5’ and 3’ loxP sites were generated by PCR using the BAC clone. Cas9, gRNA, and targeting vectors were co-injected into fertilized eggs for gene-edited mouse production and pups were genotyped by PCR followed by sequencing analysis. Mice bearing the targeted allele were backcrossed with C57BL/6J mice up to F8 generation.

Ctsk^tm1(cre)Ska^ (Ctsk^Cre^), B6N.FVB-Tg(Dmp1-cre)1Jqfe/BwdJ (Dmp1-Cre), B6.Cg-Tg(Prrx1-cre)1Cjt/J (Prrx1^Cre^), B6;SJL-Tg(Col2a1-cre)1Bhr/J (Col2-Cre), and B6.Cg-Tg(Sp7-tTA,tetO-EGFP/cre)1Amc/J (Osx1-GFP::Cre) mice have previously been reported [8, 17, 45–47]. While the Ctsk-Cre strain is a knockin mouse line, other strains (Dmp1-Cre, Prrx1-Cre, Col2-Cre, and Osx-Cre) are transgenic lines. The Gt(ROSA)26Sor^tm4(ACTB-tdTomato,-EGFP)Luo^ (Rosa26^mTmG^) mice were purchased from Jackson Laboratories. To generate experimental animals, *Chmp5^fl/fl^* mice were crossed with *Chmp5^Ctks/+^* and *Chmp5^Dmp1/+^* mice, respectively. To generate reporter mouse lines, *Chmp5^Ctks/+^* mice were mated with *Chmp5^fl/fl^*;*Rosa26^mTmG^* mice, and Ctsk^Cre^ mice were crossed with Rosa26^mTmG^ animals.

All mice were kept in C57BL/6J background and housed in the standard animal facility on a 12-hour light/dark cycle with ad libitum access to water and standard chow. In most situations, the littermates were compared otherwise as indicated. All animals were used in accordance with the Guide for Care and Use of Laboratory Animals (NIH, United States) and were handled according to protocols approved by IACUC at the University of Massachusetts Medical School (Worcester, United States) and Capital Medical University (Beijing, China). The biological and technical replicates (*n* numbers) for each animal experiment are reported in the figure legends. In this study, both sexes of mice were used and, unless otherwise stated, no gender differences were observed in the reported phenotypes.

### Micro-CT and radiography

Micro-CT analyses were performed using the Scanco μCT-35 or μCT-40 scanner (Scanco Medical). Images were processed using the Inveon Research Workplace (Siemens Medical Solutions). X-ray images were acquired using the X-Ray MX-20 Specimen (Faxitron) following the default programme.

### Histology, immunohistochemistry, and immunofluorescence

All samples were fixed in 10% neutral formalin or 4% paraformaldehyde overnight for premature samples or 2 days for mature skeletal samples. Samples were decalcified with 14% EDTA and serial sections were cut for all samples. At least five slides were selected at a certain interval (80-100 μm) from each sample for histological evaluation. Fate-mapping images were acquired using a Leica TCS SP5 II laser scanning confocal microscope.

Immunofluorescence for γH2AX was performed using a Phospho-Histone H2A.X (Ser139) antibody (clone 20E3, CST). Immunohistochemistry for Ki-67 was performed using the VECTASTAIN® Elite® ABC HRP Kit (Peroxidase, Rabbit IgG; Vector Laboratories). Cells for immunofluorescence were cultured in 35 mm MatTek glass bottom dishes (No. 1.5 coverslip, MatTek Corporation). Briefly, tissues or cells were fixed in 4% PFA for 15 min, blocked in 1% BSA for 30 min, and incubated with primary antibodies for 2 hrs and then fluorescence conjugated secondary antibodies for 1 hr. Images were acquired using the Leica TCS SP5 II laser scanning confocal microscope. The quantification of intracellular vesicles was performed using FIJI software following the particle analysis procedure (https://imagej.net/imagej-wiki-static/Particle_Analysis, NIH).

### Bone mechanical test

The femurs of *Chmp5^Ctsk^* and *Chmp5^fl/fl^* mice at 6-7 weeks of age were subjected to mechanical testing using the DMA Q800 mechanical test system with the 3-point bending mode following the manufacturer’s procedure (TA Instruments, USA). The femurs were placed in a direction of the anterior surface downward and the posterior aspect of the femoral condyles facing upward. The span length (L) was 10 mm and the load was applied in the middle of the bone. Five mice per group were used and bone stiffness and ultimate stress (fracture stress) from male mice were reported. Similar changes were observed in both sexes.

### Tests of skeletal muscle functions

The four-limb hanging test was performed using the top grid of a mouse cage [48]. The grid was set at a height of about 35 cm, and the measurement was repeated for 3-5 times for each mouse at a rest interval of about 3-5 min. The maximum hang time was recorded for each animal. Both male and female animals were included in this experiment, and there was no significant sex difference regarding the four-limb hanging time.

The forelimb grip strength test was carried out using an electronic scale as described with slight modification [48, 49]. A mouse was allowed to grasp the gauze attached on the scale. The scale was reset to 0 g after stabilization and a researcher slowly pulled the mouse’s tail backward. Five consecutive measurements were performed for each mouse and the peak pull force in grams was recorded. A significant sex difference was observed in the forelimb grip strength. Both male and female mice were included in this study and the same trend of changes was found in both genders.

### Skeletal progenitor cell culture and sorting

The periskeletal progenitors were isolated from the hindlimbs of *Ctsk-Cre;Rosa26^mTmG/+^*, *Chmp5^Ctsk/+^;Rosa26^mTmG/+^*, and *Chmp5^Ctsk^;Rosa26^mTmG/+^* mice (*n* = 10 animals per genotype) at 2 weeks of age according to the previously established method [23]. Briefly, the periskeletal tissues were dissected, minced into small pieces, and digested with 1 mg/ml collagenase type I (Worthington), 1 mg/ml collagenase type II (Worthington) and 1.5 mg/ml dispase (Roche) for 30 min at 37°C. Cells were cultured in skeletal stem cell medium (Joint Therapeutics, Beijing, China) for one week, stained with anti-CD31 and anti-CD45 antibodies, and sorted using a FACSAria II cell sorter (BD Biosciences) for further analyses.

In all experiments, *Chmp5^Ctsk^* and control cells from littermate mice were cultured, sorted, and re-seeded in parallel for subsequent analyses. The coculture experiment was carried out by directly mixing 90% *wild-type* skeletal progenitors with 10% *Chmp5^Ctsk^* or control periskeletal progenitors to simulate the in vivo context of cell-cell contact in periskeletal overgrowth. The cocultures were induced in osteogenic differentiation medium for 4 weeks. The biological and technical replicates (*n*-numbers) for each cellular experiment are reported in the figure legends.

### CRISPR/CAS9 lentiviral infection

Pairs of CRISPR guide RNA oligos (mouse *Chmp5* single guide RNAs [sgRNAs] targeting GGCTCCGCCACCTAGCTTGA and GTTTCGCTTTTCCGAAGAAT on exon 1 respectively) were annealed and cloned into the BsmBI sites of the lentiCRISPR V2-puro vector (plasmid 52961, Addgene). CRISPR lentiviral plasmids and lentiviral packaging plasmids (pMDLg/pRRE, pRSV-Rev, and pMD2.G; Addgene) were transfected into HEK 293T cells. The supernatants were harvested and filtered through a 0.45-μm filter 2.5 days after transfection. MC3T3-E1 and ATDC5 cells were infected with CRISPR lentivirus and selected with puromycin (4.5 μg/ml, Clontech) for 7 days. The depletion of *Chmp5* was confirmed by Western blotting.

### *In vitro* osteogenesis

For osteogenic differentiation, 1.0 x 10^5^ cells were seeded in each well of 24-well plates and cultured overnight. Cells were changed into osteogenic media (Joint Therapeutics) the next day and induced for the indicated times. Osteogenesis was determined by alizarin red and von Kossa staining and by examining the activity of alkaline phosphatase.

### Cell cycle, proliferation, and apoptosis analyses

The cell number was counted with a Countess II FL Automated Cell Counter (ThermoFisher Scientific, USA) at indicated time intervals. The AlamarBlue cell viability assay was performed as previously described [23]. The cell cycle was determined using the APC-BrdU Flow Kit (BD Biosciences). In vitro apoptosis analysis was carried out with Annexin V-PI staining and in situ cell apoptosis was determined with TUNEL staining (Abcam).

### RNA-seq, data analysis, and functional annotation

The sorted *Chmp5^Ctsk^* and *Chmp5^Ctsk/+^* periskeletal progenitors were subjected to RNA-seq For library preparation, 100 ng of total RNA from each sample was subjected to rRNA depletion with the NEBNext rRNA Depletion Kit (New England Biolabs) according to the manufacturer’s manual. Subsequently, the rRNA-depleted RNA was used to build the RNA-seq library with the NEBNext Ultra II Directional RNA Library Prep Kit (New England Biolabs) according to the manufacturer’s manual. The index of each RNA-seq library was introduced using NEBNext Multiplex Oligos for Illumina (New England Biolabs). Each RNA-seq library was analyzed by the fragment analyzer (Advanced Analytical Technologies) and quantified by the KAPA Library Quantification Kit (KAPA Biosystems) according to the manufacturer’s manual. An equal amount of each RNA-seq library was mixed for sequencing in one lane of Illumnia HiSeq3000 in paired-end 150 base mode (Illumina).

The raw Illumina pair-end reads were first aligned with ribosomal RNA (GenBank ID BK000964.1) with Bowtie2 v2.2.6 [50]. Reads that failed to map to ribosomal RNA were aligned with the mouse reference genome mm10 with STAR v2.5.3 [51]. Differential analysis was performed with Cuffdiff v2.2.1 [52]. The raw data were analyzed by two independent statisticians and similar results were obtained.

Pathway analyses were performed for differentially expressed genes using the Reactome online analysis tool [53] and the Ingenuity Pathway Analysis software (Qiagen). Gene ontology term over-representation analysis was run using PANTHER (version 14.1) with all genes expressed in a tissue or cell type as background and the differentially expressed gene list as input. The P-values of Fisher’s exact tests were corrected by controlling the false discovery rate (FDR). Gene set enrichment analyses were performed using GSEA or ConsensusPathDB. For GSEA-based analysis, gene sets of interests were downloaded from the MsigDB, while all built-in gene sets were used for ConsensusPathDB-based analysis.

### Nano LC-MS/MS for secretomic and proteomic analyses

Label-free relative protein quantification analyses were performed to identify differentially expressed proteins in the supernatant or cell lysis of *Chmp5^Ctsk^* versus control periskeletal progenitors (*n* =3 animals per group). For cell supernatants, the same amounts of samples went through 3 kD ultrafiltration tubes and then were processed for reduction and alkylation. Subsequently, the samples were digested in trypsin overnight and the peptides were desalted prior to mass spectrometry analysis. For cell lysis, samples were precipitated using acetone after reduction and alkylation. The samples were then subjected to digestion, desalination, and mass spectrometry analysis.

Quantitative data were collected on an Orbitrap Eclipse™ Tribrid^TM^ mass spectrometer (ThermoFisher Scientific) coupled to a nanoLC system (EASY-nLC™ 1200, ThermoFisher Scientific). All peptide mixtures were analyzed using the same chromatographic conditions: 5 μl of each sample were fractionated in a 150 μm×15 cm in-house made column packed with Acclaim PepMap RPLC C18 (1.9 μm, 100 Å, Dr. Maisch GmbH, Germany) working at a flow rate of 600 nl/min, using a linear gradient of eluent B (20% 0.1% formic acid in water + 80% acetonitrile) in A (0.1% formic acid in MilliQ water) from 4% to 10% for 5 min, 10% to 22% for 80 min, 22% to 40% for 25 min, 40% to 95% for 5 min, and 95% to 95% for 5 min. MS/MS analyses were performed using Data-Dependent Acquisition (DDA) mode: one MS scan (mass range from 350 to 1500 m/z) was followed by MS/MS scans up to the top 20 most intense peptide ions from the preview scan in the Orbitrap, applying a dynamic exclusion window of 120 seconds.

The raw MS files were analyzed and searched against the Mus_musculus_reviewed protein database using MaxQuant (1.6.2.10). The parameters were set as follows: the protein modifications were carbamidomethylation (C) (fixed), oxidation (M) (variable), acetyl (Protein N-term) (variable); the enzyme specificity was set to trypsin; the maximum missed cleavages were set at 2; the precursor ion mass tolerance was set at 20 ppm; and the MS/MS tolerance was 20 ppm. Only highly confident identified peptides were chosen for downstream protein identification analysis.

### Western blotting

Western blot was performed using 4–20% Mini-PROTEAN® TGX™ precast protein gels (Bio-Rad). The following antibodies: anti-CHMP5 polyclonal antibody [6], anti-COL1A1 monoclonal antibody (clone E8F4L, Cell Signaling Technology, CST), anti- AGLN polyclonal antibody (cat#: AB14106, Abcam), anti-p16 INK4A monoclonal antibody (clone E5F3Y, CST), anti-p21 Waf1/Cip1 monoclonal antibody (clone E2R7A, CST), anti-VPS4A monoclonal antibody (clone A-11, Santa Cruz Technology), total OXPHOS antibody cocktail (Abcam), anti-GAPDH monoclonal antibody (clone 14C10, CST), and anti-β-Actin monoclonal antibody (clone D6A8, CST), were used.

### Quantitative PCR

Total RNA was isolated from cells using Trizol reagent (Qiagen). RNA samples were treated with the TURBO DNA-free^TM^ Kit (Thermo Fisher) and equal amounts (1–2 μg) were used for reverse transcriptase reaction using an iScript™ Reverse Transcription Supermix (Bio-Rad). Quantitative PCR was run using the SYBR® Green Master Mix on a CFX Connect™ Real-Time PCR Detection System (Bio-Rad). Gene expression levels were analyzed relative to the housekeeping gene *Gapdh* and presented by 2^−△Ct^ [54]. The primer sequences were used for qPCR are as following: *Chmp5,* forward, 5’-ATGAGAGAGGGTCCTGCTAAG-3’, reverse, 5’-CCGTGGTCTTGGTGTCCTTTA-3’; *H2afx*, forward, 5’-GTGGTCTCTCAGCGTTGTTC-3’, reverse, 5’-CGGCC TACAGGGAACTGAA-3’; *Tp53*, forward, 5’-GTCACAGCACATGACGGAGG-3’, reverse, 5’-TCTTCCAGATGCTCGGGATAC-3’; *Gapdh*, forward 5′-TGCCAGCCTCGTCCCG TAGAC-3′, reverse 5′-CCTCACCCCATTTGATGTTAG-3′; *VPS4A*, forward 5′-AGAACCAGAGTGAGGGCAAGGG-3′, reverse 5′-GCACCCATCAGCTGTTCTTGCA-3′; *CHMP5*, forward 5′-CCAGCCTGACTGACTGCATTGG-3′, reverse 5′-TCGCAAGGCTTTCTGCTTGACC-3′; *GAPDH*, forward 5′-GGAGTCCACTGGCGTCTTCAC-3′, reverse 5′-GAGGCATTGCTGATGATCTTGAGG-3′.

### Extracellular vesicle tracking

The concentration and size distribution of extracellular vesicles were analyzed using the NanoSight NS300 following the manufacturer’s protocol (Malvern Instruments). Briefly, cells were seeded in 24-well plates at the density of 1 x 10^5^ cells per well and cultured overnight. The next morning, cells were changed to serum-free medium and incubated for 9 hrs. The medium was collected and subjected to sequential centrifugation at 300 g for 10 min, 2000 g for 20 min, and 10000 g for 30 min. The resulting supernatant was manually injected into the instrument and run in the “standard measurement” module with five captures per sample and the data were processed with a detection threshold of 2.

### Live-cell imaging for cell endocytosis

The *wild-type* and *Chmp5^Ctsk^* periskeletal progenitors were incubated on ice for 10 min before adding fresh medium with 2 ug/ml pHrodo™ red EGF conjugate (ThermoFisher). Cells were incubated at 37 °C for 30 min and subsequently washed and changed to fresh medium. Live-cell images were obtained at 1, 12, and 24 hrs using the Leica TCS SP5 II laser scanning confocal microscope.

### Transmission electron microscopy

*Wild-type* and *Chmp5^Ctsk^* periskeletal progenitors were fixed in culture plates overnight at 4 °C using 2.5% glutaraldehyde in 0.1 M Na cacodylate-HCl buffer (pH 7.2). After washing in 0.5 M Na cacodylate-HCl buffer (pH 7.0), the cells were post-fixed for 1 hr in 1% osmium tetroxide (w/v) at room temperature. After post-fixation, the culture plate with adherent cells were enblock stained (20 min) with 1% aqueous uranyl-acetate (w/v). Culture plates were washed again in the same buffer and dehydrated through a series of graded ethanol to 100% and transferred through two changes of 50/50 (v/v) SPIpon resin (Structure Probe, Inc.)/100% ethanol and left overnight to infiltrate. The following morning, the cell culture plates were transferred through three changes of fresh SPIpon resin to finish the infiltration and embedding and finally, the plates were filled with freshly prepared SPIpon resin and polymerized for 48 hrs at 70 °C.

Once fully polymerized, the plate was cut apart, and each well was plunged into liquid nitrogen to separate the SPIpon epoxy block with the embedded cells from the culture dish. The round epoxy disks with the embedded cells were then examined under an upright light microscope, and areas of cells were cut from the disks and glued onto blank microtome studs and trimmed for ultramicrotomy. Ultrathin sections (70 nm) were cut on a Reichart-Jung ultramicrotome using a diamond knife. The sections were collected and mounted on copper support grids and contrasted with lead citrate and uranyl acetate and examined on a FEI Tecnai G2 Spirit transmission electron microscope at 100 Kv accelerating voltage and images were recorded at various magnifications using a Gatan 2K digital camera system.

### Mitochondrial respiration

Mitochondrial respiration was examined in *Chmp5^Ctsk^* and *wild-type* periskeletal progenitors using a Seahorse XF^e^96 Extracellular Flux Analyzer (Agilent) as previously described [55]. Briefly, 10000 cells per well were seeded in XF 96-well culture plates and cultured overnight. The next day, cells were changed to assay medium and incubated at 37 °C w/o CO2 for 1 hr. The assay was carried out using the Seahorse XF Cell Mito Stress Test kit with sequential injections of Oligomycin (1 µM), FCCP (1 µM), and Rotenone plus Antimycin A (0.5 µM) following the manufacturer’s protocol.

### Senolytic treatments

For in vitro cell treatment, *Chmp5^Ctsk^* and control cells were seeded in 96-well plates at a density of 6000 cells/ well and cultured overnight. Quercetin and dasatinib were added to the culture medium at a final concentration of 50 μM and 500 nM, respectively. Apoptotic cells are labeled using Incucyte Annexin V Red Dye and are quantified in real time using the Incucyte Live-Cell Analysis System. Six replicates were used for each group and the experiment was repeated twice using cells from two different animals per group.

For in vivo treatment, the senolytic drugs quercetin and dasatinib were administered intraperitoneally weekly at 50 µg/g and 5 µg/g body weight, respectively. Senolytic treatment in *Chmp5^Ctsk^* and littermate control mice started from the first week after birth and mice were collected at 7-8 weeks of age for skeletal analyses. For the *Chmp5^Dmp1^* strain, treatment began the second week after birth and the animals were monitored for 16 weeks. ABT-263 was administered intraperitoneally to *Chmp5^Ctsk^* and littermate control mice weekly at 10 ug/g body weight, starting from the first week after birth and mice were collected at 8 weeks of age for micro-CT analyses.

### Statistics

All data are represented as mean ± s.d.. Normality and equal variance of the data sets were tested using the Shapiro-Wilk test and the F test, respectively. The data were determined to be normally distributed and have equal variance unless otherwise specified. For experiments with three or more groups, statistical analysis was performed using one-way or two-way ANOVA followed by multiple comparisons. For comparisons between two groups, the two-tailed unpaired Student’s *t*-test was applied. All analyses were performed with Prism 10.1.1 (GraphPad).

## Supplementary Figure Legends

**Fig. S1.**
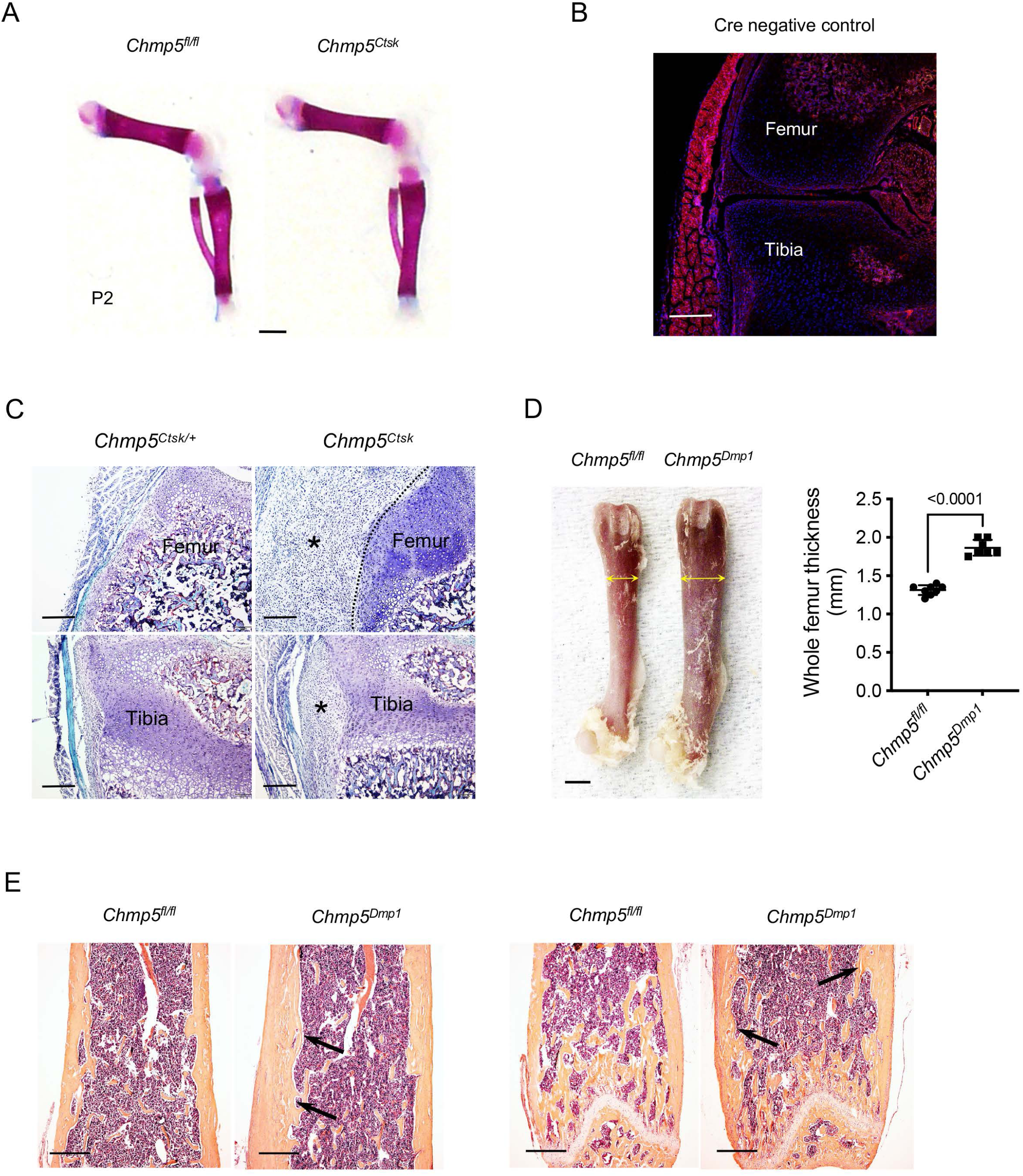
CHMP5 restricts bone formation in osteogenic lineage cells. (**A**) Alizarin red staining of skeletal preparations on postnatal day 2. *n* = 4 mice per group. Scale bar, 1 mm. (**B**) Confocal images of *Rosa26^mTmG/+^* (Cre^−^) control mice. Related to Fig. 1E. Scale bar, 200 μm. (**C**) Tartrate-resistant acid phosphatase (TRAP) staining demonstrating the absence of osteoclasts in peri-skeletal overgrowth (asterisk) of *Chmp5^Ctsk^* mice at 2 weeks of age. TRAP^+^ osteoclasts in the bone marrow as positive control; dot-line representing approximate border between peri-skeletal overgrowth and the bone at femoral condyle of the knee. Images are representative of 3 animals per group. Scale bar, 200 μm. (**D**) Gross image and measurement of femur thickness showing bone expansion in *Chmp5^Dmp1^* versus *Chmp5^fl/fl^* mice at 10 weeks of age. *n* = 13 *Chmp5^fl/fl^* and 12 *Chmp5^Dmp1^* mice. Scale bar, 1 mm. (**E**) H&E histology showing bone expansion at the endosteum (arrows) in *Chmp5^Dmp1^* mice. The left images showing the midshaft and the right images showing the metaphysis of the femur bone. Images are representative of 3 mice per group. Scale bars, 0.5 mm. Data are mean ± s.d.; two-tailed unpaired Student’s *t*-test.

**Fig. S2.**
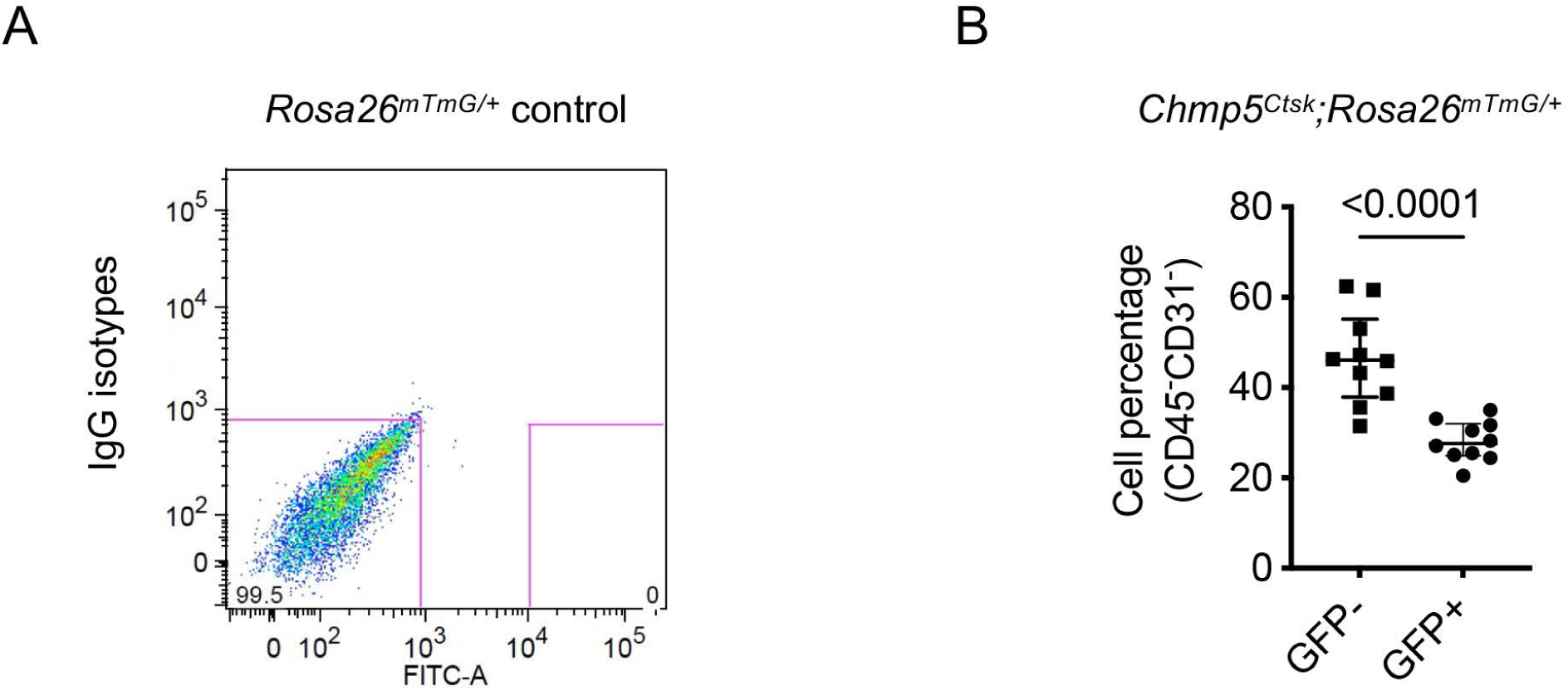
Sorting of CD45^−^;CD31^−^;CTSK^+^ periskeletal progenitors. (**A**) Unstained cells from *Rosa26-mTmG^fl/+^* mice as a negative control for sorting CD45^−^;CD31^−^;CTSK^+^ skeletal progenitors. Related to Fig. 2A. (**B**) Quantification of CD45^−^;CD31^−^;GFP^+^ and CD45^−^;CD31^−^;GFP-cell population in samples of *Chmp5^Ctsk^;Rosa26^mTmG/+^* mice. *n* = 10 mice per group.

**Fig. S3.**
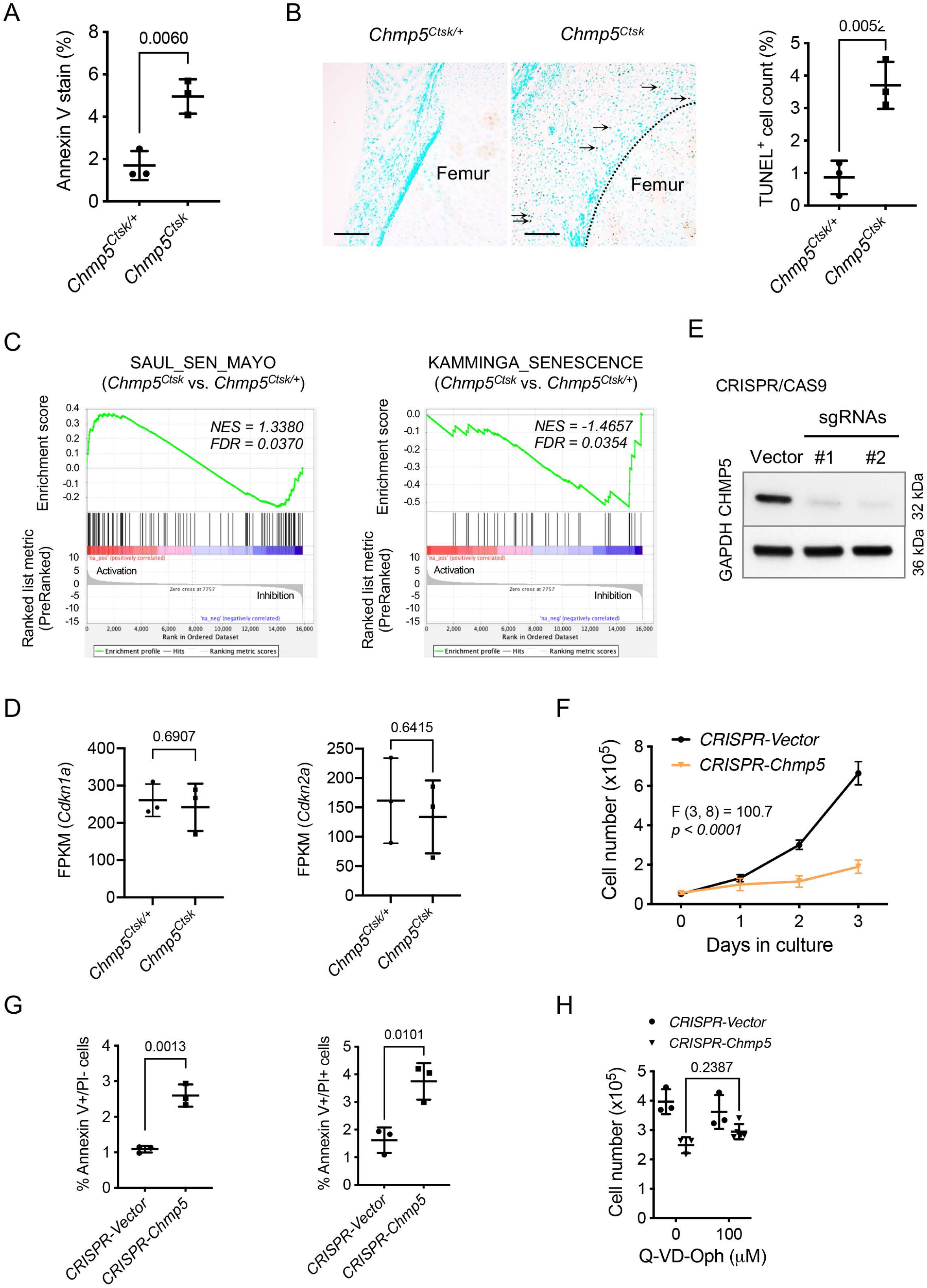
Cell apoptosis is not the main cause of the decreased proliferative rate in *Chmp5*-deficient skeletal progenitors. (**A**) Annexin V stain examining cellular apoptosis in *Chmp5^Ctsk^* and *Chmp5^Ctsk/+^* periskeletal progenitors. *n* = 3 replicates each group; repeated twice using cells from 2 mice per group. (**B**) TUNEL staining and quantification of TUNEL^+^ cells (%) in the periskeletal tissues of *Chmp5^Ctsk^* and *Chmp5^Ctsk/+^* mice. Arrows indicating positive cells in periskeletal overgrowth in *Chmp5^Ctsk^* mice, dot-line representing approximate border between periskeletal overgrowth and bone in the femoral condyle of the knee; *n* = 3 animals per group; scale bars, 100 μm. (**C**) GSEA of RNA-seq data showing significant enrichment of SAUL_SEN_MAYO and KAMMINGA_SENESCENCE genesets in *Chmp5^Ctsk^* relative to *Chmp5^Ctsk/+^* periskeletal progenitors. (**D**) *Cdkn1a* and *Cdkn2a* mRNA levels in *Chmp5^Ctsk^* and *Chmp5^Ctsk/+^* periskeletal progenitors according to RNA-seq data. *n* = 3 per group with cells from 3 different animals. (**E**) Western blot confirming deletion of *Chmp5* in ATDC5 cells by lentiviral CRISPR/CAS9. (**F**) Cell number counting determining cell proliferation in ATDC5 cells after deleting *Chmp5* by lentiviral CRISPR/CAS9. *n* = 4 replicates per group per time point, experiment repeated three times. (**G**) Annexin V-PI stain analyzing cellular apoptosis in ATDC5 cells with or without *Chmp5* deletion. *n* = 3 replicates for each group, repeated 3 times. (**H**) Cell number counting determining cell proliferation in ATDC5 cells with or without *Chmp5* deletion after treatment with the pan-caspase inhibitor Q-VD-OPh. *n* = 3 replicates of each group each dose, repeated twice. All data are mean ± s.d.; two-tailed Student’s *t*-test for comparison of two groups or 2-way ANOVA followed by multiple comparisons.

**Fig. S4.**
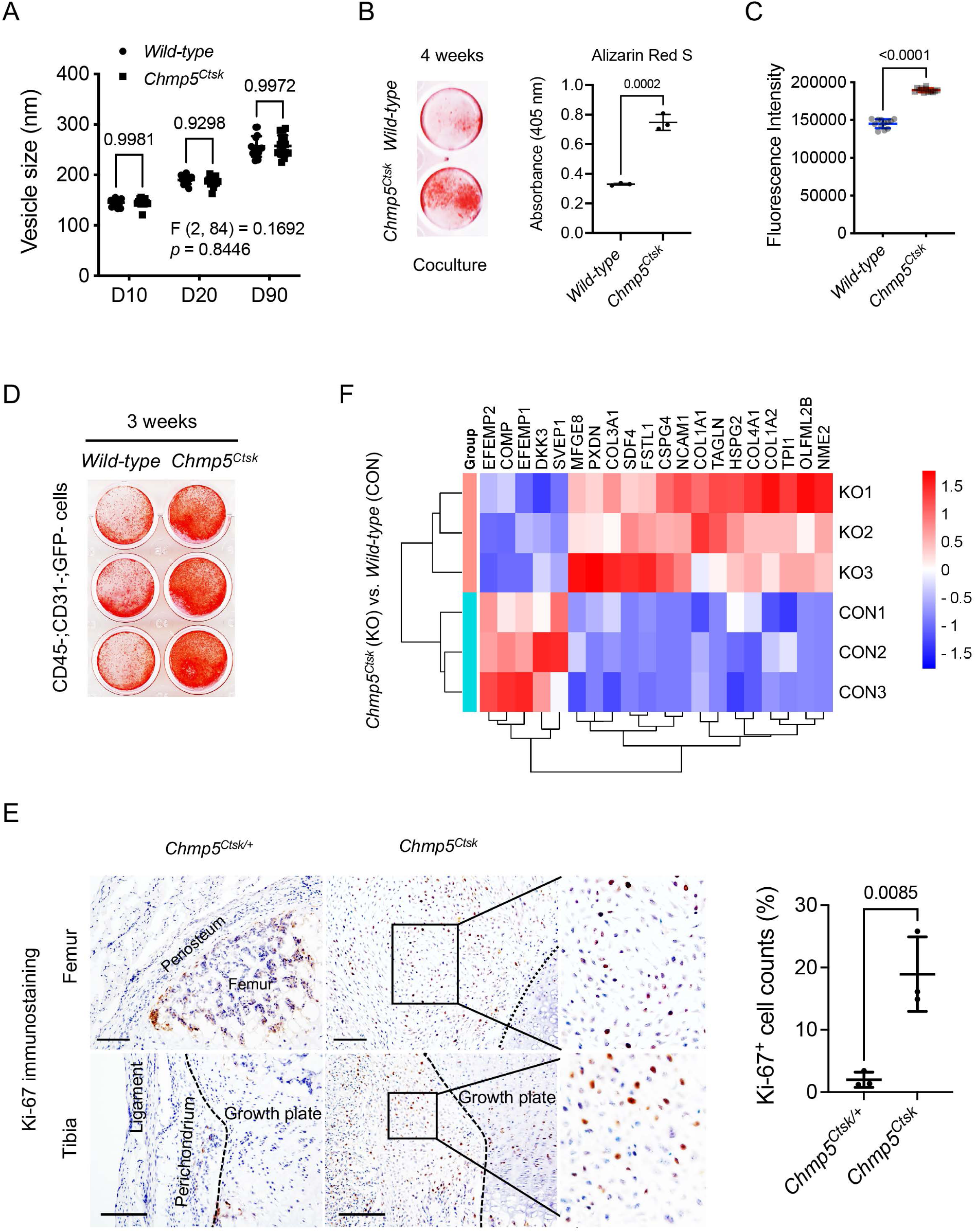
Secretory phenotype of *Chmp5*-deficient skeletal progenitors. (**A**) Nanoparticle tracking analysis showing the size distribution of extracellular vesicles in *Chmp5^Ctsk^* and *wild-type* skeletal progenitors. Data pooled from 3 replicates, 5 reads for each; repeated twice using cells from 2 mice; D10, D50, and D90 indicating percent undersize, for example D50 = 229 nm representing that 50% of vesicles are 229 nm or smaller. (**B**) Alizarin red staining and quantification showing osteogenesis in coculture of *Chmp5^Ctsk^* and *wild-type* periskeletal progenitors. *n* = 3 replicates per group, repeated twice using cells from 2 different animals. (**C**) AlarmaBlue assay examining cell proliferation in neighboring CD45^−^;CD31^−^;GFP-progenitors sorted from periskeletal tissues of *Chmp5^Ctsk^;Rosa26^mTmG/+^* or *Ctsk-Cre;Rosa26^mTmG/+^* mice. *n* = 12 replicates per group per time-point, repeated 3 times using cells from 3 animals. (**D**) Alizarin red staining determining osteogenesis in neighboring CD45^−^;CD31^−^;GFP-progenitors sorted from periskeletal tissues of *Chmp5^Ctsk^;Rosa26^mTmG/+^* or *Ctsk-Cre;Rosa26^mTmG/+^* mice after induction in osteogenic medium for 3 weeks. *n* = 3 replicates per group per time point, repeated 3 times using cells from 3 animals. (**E**) Immunostaining and quantification of cell proliferation marker Ki-67 in the periskeletal tissues around the knee of *Chmp5^Ctsk^* and *Chmp5^Ctsk/+^* mice. *n* = 3 animals per group; dot-line representing the approximate border between the periskeletal overgrowth and the bone in the femoral condyle; scale bars, 100 μm. (**F**) Heatmap of LC-MS/MS results showing differentially secreted proteins in supernatants of *Chmp5^Ctsk^* relative to *wild-type* skeletal progenitors. *n* = 3 with cells from 3 different animals per group for Nano LC-MS/MS analysis. All data are mean ± s.d., 2-way ANOVA followed by multiple comparisons or two-tailed unpaired Student’s *t*-test for 2-group comparison.

**Fig. S5.**
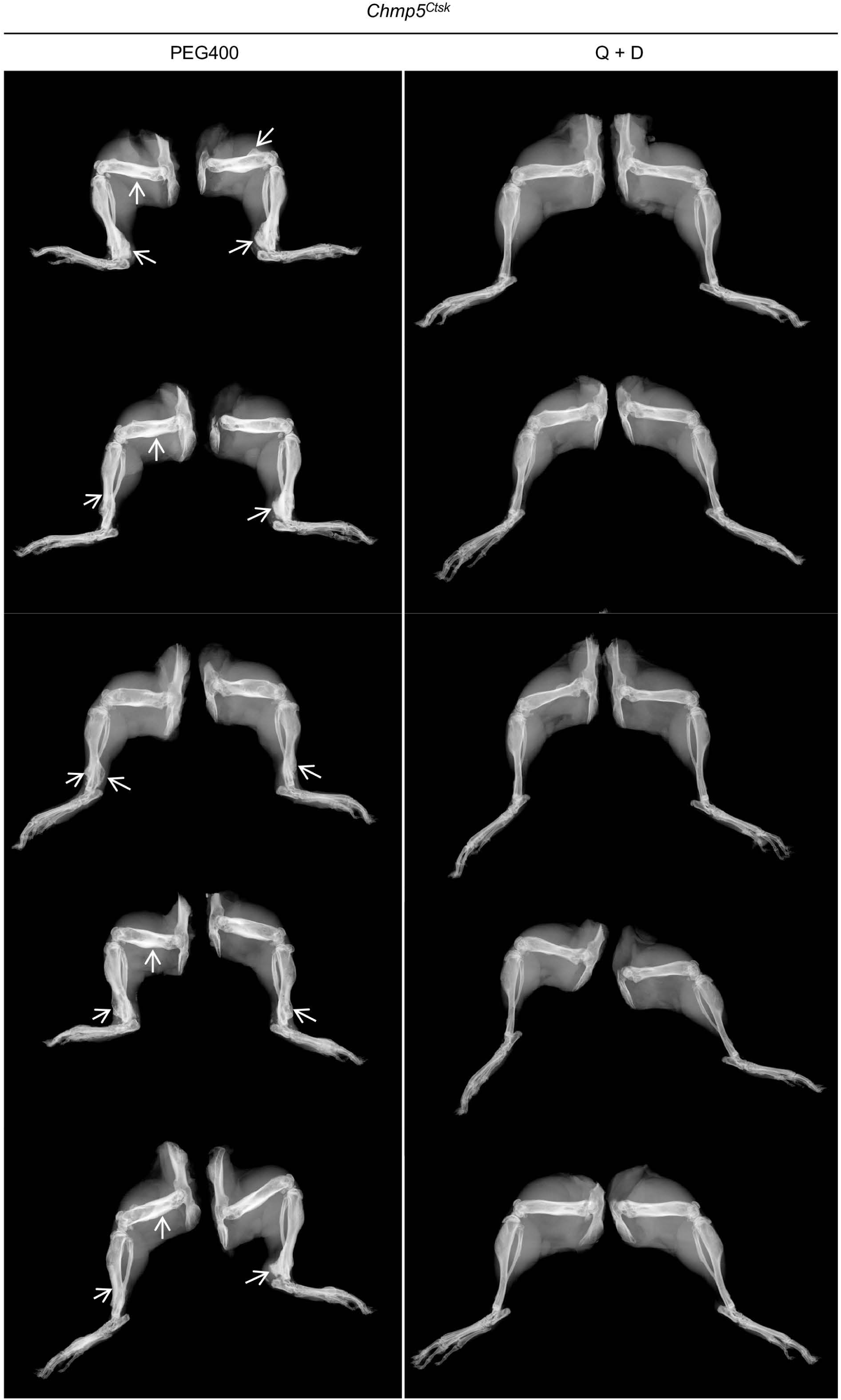
Additional x-ray images of *Chmp5^Ctsk^* mice after treatment with Q + D or the vehicle PEG400 for 7 weeks. Arrows indicating periskeletal bone overgrowths.

**Fig. S6.**
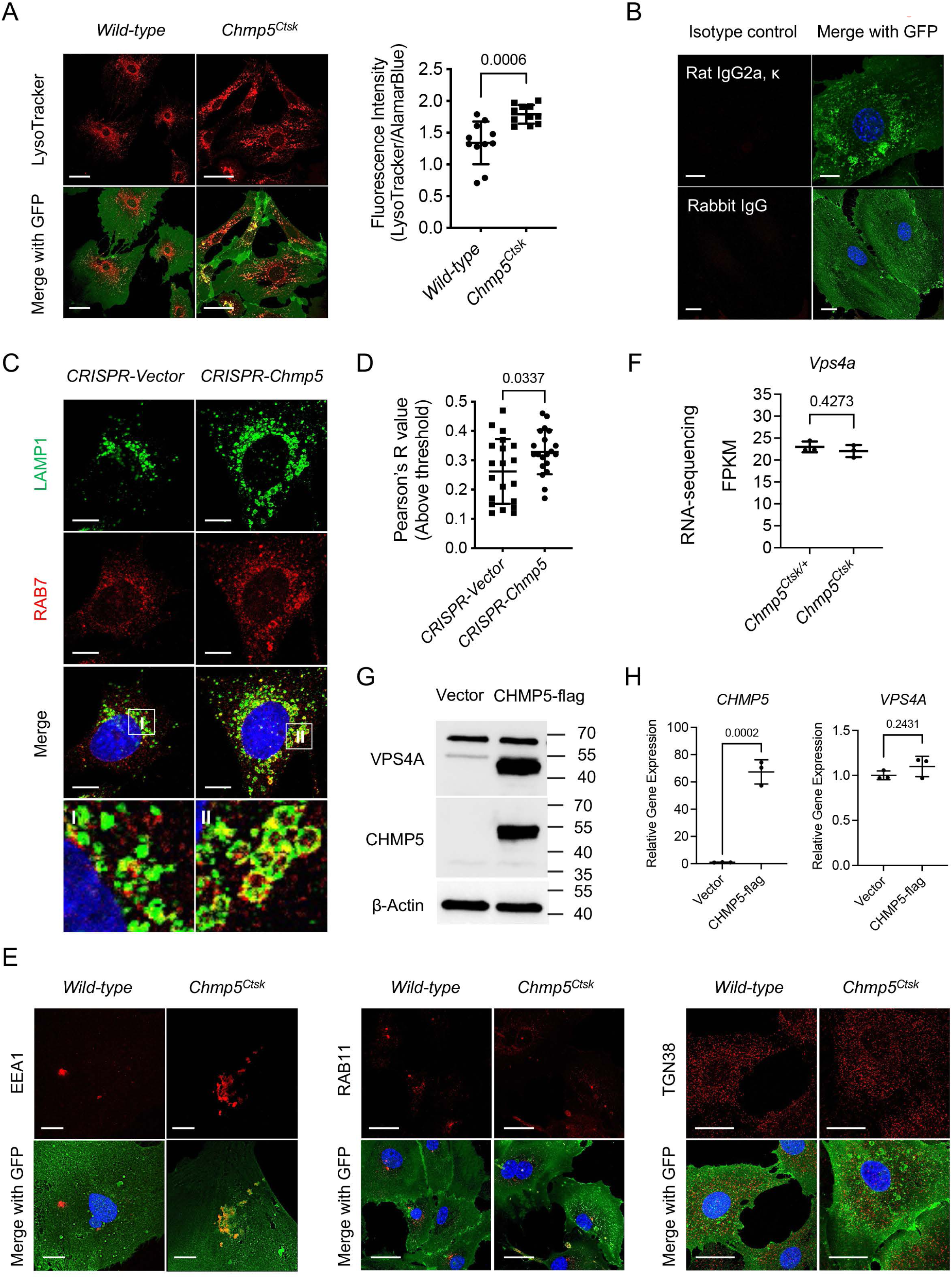
CHMP5 is essential for endolysosomal functions and maintaining VPS4A protein in skeletal progenitors. (**A**) Representative confocal fluorescence images and quantification of the fluorescence intensity of LysoTracker Red DND-99 in *Chmp5^Ctsk^* compared to *wild-type* periskeletal progenitors. *n* = 11 replicates per group for quantitative analysis; repeated 3 times using cells from 3 mice. Scale bars, 50 μm. (**B**) Isotype controls for immunofluorescence staining. Scale bars, 20 μm. (**C**) Co-localization of LAMP1 and RAB7 in *Chmp5*-sufficient or *Chmp5*-deficient ATDC5 cells. Scale bars, 10 μm. (**D**) Colocalization analysis of LAMP1 and RAB7 performed by the ImageJ Coloc2 programme and Pearson’s R value (above threshold) shown, *n* = 20 cells per group. (**E**) Representative confocal images of immunostaining for early endosome marker EEA1, cycling endosome marker RAB11, and trans-Golgi network marker TGN38 in *Chmp5^Ctsk^* vs. *wild-type* periskeletal progenitors. *n* = 30 cells per group. Scale bars, 10 μm (EEA1), 20 μm (RAB11), 25 μm (TGN38). (**F**) RNA-seq data showing *Vps4a* mRNA expression in *Chmp5^Ctsk^* vs. *Chmp5^Ctsk/+^* periskeletal progenitors. *n* = 3 with cells from 3 different animals for each group. (**G**) Western blot showing that CHMP5 and VPS4A protein expression after transfection of CHMP5 or empty vector in HEK-293T cells. Experiments repeated 3 times with independent cells and transfections at different times. (**H**) Quantitative PCR showing the expression of *CHMP5* and *VPS4A* mRNA after transfecting CHMP5 or empty vector in HEK-293T cells. *n* = 3 replicates for each group, experiments repeated 3 times with independent cells and transfections at different times. All data shown as mean ± s.d.; two-tailed unpaired Student’s *t*-test.

**Fig. S7.**
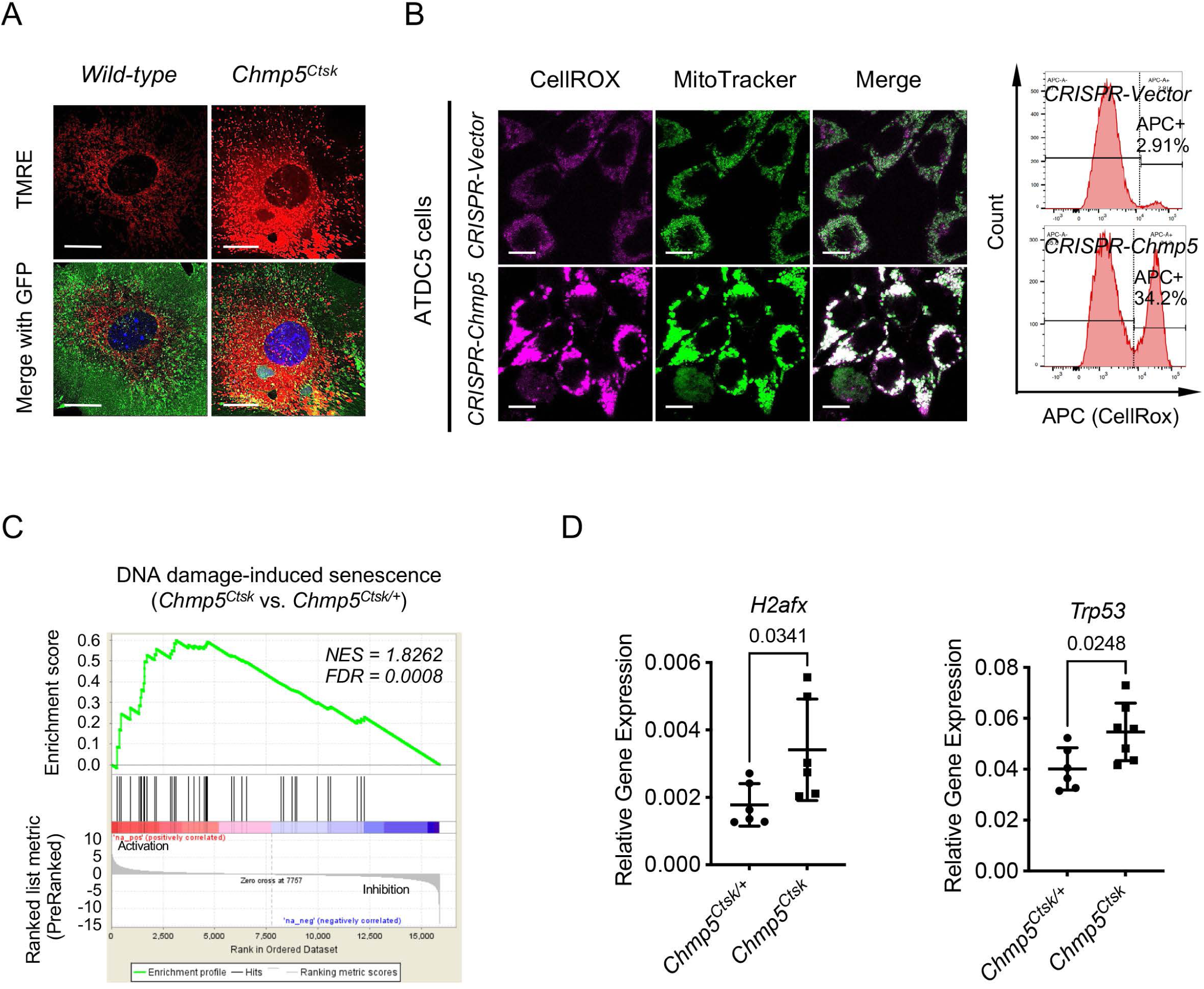
Mitochondrial dysfunction and activation of DNA damage-induced senescence in *Chmp5*-deficient skeletal progenitors. (**A**) Representative TMRE confocal images showing accumulation of mitochondria in *Chmp5^Ctsk^* periskeletal progenitors. *n* = 30 cells per group, scale bars, 25 μm. (**B**) Confocal fluorescence imaging mapping intracellular ROS (CellROX Deep Red) and mitochondria (MitoTracker Green) in ATDC5 cells with or without *Chmp5* depletion. *n* = 20 cells for each group; scale bars, 10 μm. The right histogram showing the quantification of CellROX Deep Red in ATDC5 cells by flow cytometry. *n* = 3 per group, experiment repeated twice. (**C**) GSEA of RNA-seq data showing positive enrichment of genes associated with the molecular pathway of DNA damage-induced senescence in *Chmp5^Ctsk^* relative to *Chmp5^Ctsk/+^* periskeletal progenitors. (**D**) Quantitative PCR determining expression of *H2afx* and *Trp53* genes in *Chmp5^Ctsk^* relative to *Chmp5^Ctsk/+^* periskeletal progenitors. *n* = 6 with cells from different animals per group. Data shown as mean ± s.d.; two-tailed unpaired Student’s *t*-test.

